# Human complement Factor H and Properdin act as soluble pattern recognition receptors and differentially modulate SARS-CoV-2 Infection

**DOI:** 10.1101/2023.07.07.548083

**Authors:** Nazar Beirag, Praveen M Varghese, Chandan Kumar, Susan Idicula-Thomas, Martin Mayora Neto, Haseeb A. Khan, Robert B. Sim, Taruna Madan, Nigel Temperton, Uday Kishore

## Abstract

Severe cases of SARS-CoV-2 infection are characterised by an imbalanced immune response, excessive inflammation, and the development of acute respiratory distress syndrome, which can lead to multiorgan failure and death. Several studies have demonstrated dysregulated complement activity as an indicator of immunopathogenesis in the SARS-CoV-2 infection. Notably, the complement alternative pathway has been implicated in driving the excessive inflammation during severe SARS-CoV-2 infection. Reduced levels of factor H (FH), a down-regulator of the alternative pathway, and increased levels of properdin (Factor P/FP), the only known up-regulator of the alternative pathway, have been observed in individuals with severe COVID-19 infection. The present study investigated the complement activation-independent, and a more direct role of FH and FP against SARS-CoV-2 infection. Using direct ELISA, the interactions of FH and FP with the SARS-CoV-2 spike (S) and receptor binding domain (RBD) were assessed. Using S protein expressing lentiviral pseudotypes, the cell binding and luciferase-based virus entry assays were employed to assess the potential modulatory effects of FH, FP, and recombinant thrombospondin repeats 4 and 5 (TSR4+5) on SARS-CoV-2 cell entry. We also evaluated the immunomodulatory functions of FH and FP in the cytokine response triggered by SARS-CoV-2 pseudotypes via RT-qPCR. SARS-CoV-2 S and RBD proteins were found to bind both FH and FP. Treatment of A549 cells expressing human ACE2 and TMPRSS2 with FP or TSR4+5 resulted in increased cell entry and binding of SARS-CoV-2 pseudotypes. In silico studies revealed that FP increases affinity between SARS-CoV-2 and host ACE2. The impact of FP on viral cell entry and binding was reversed by anti-FP antibody treatment in A549-hACE2+TMPRSS2 cells. However, FH treatment reduced the cell entry and binding of SARS-CoV-2 lentiviral pseudotypes. Furthermore, the A549-hACE2+TMPRSS2 cells challenged with SARS-CoV-2 spike, envelope, nucleoprotein, and membrane protein expressing alphaviral pseudotypes pre-treated with FP or TSR4+5, exhibited upregulation of the transcripts of pro-inflammatory cytokines, such as IL-1β, IL-8, IL-6, TNF-α, IFN-α and RANTES (as well as NF-κB). Conversely, FH pre-treatment downregulated the expression of these pro-inflammatory cytokines. Treatment of A549-hACE2+TMPRSS2 cells with FP increased S protein-mediated NF-κB activation, while FH treatment reduced it. These findings suggest that FH may act as an inhibitor of SARS-CoV-2 cell entry and binding, thereby attenuating the infection-associated inflammatory response in a complement activation-independent manner. FP may contribute to viral cell entry, binding, and exacerbating the immune response. That may result in potentially influencing the severity of the infection.

## Introduction

The COVID-19 pandemic, caused by severe acute respiratory syndrome coronavirus type 2 (SARS-CoV-2), has had a significant global impact, with millions of infections and fatalities reported (1). The pathogenesis of COVID-19 is primarily characterised by dysregulation of the immune response (2). SARS-CoV-2 is an enveloped RNA virus of the Coronaviridae family (1). It consists of several structural proteins, including the nucleocapsid (N), membrane spike (S), membrane (M), and envelope (E) proteins, along with other auxiliary proteins that aid in viral entry and replication (1). The surface of SARS-CoV-2 is covered by the S protein, which comprises two subunits, S1 and S2, responsible for binding to host cell receptors (1). The virus infects various cell types, including alveolar macrophages and epithelial cells, by attaching to the angiotensin-converting enzyme 2 (ACE2) receptor (1).

The complement system plays a crucial role in both innate and acquired immunity, providing a robust defense against various infections caused by viruses, bacteria, fungi, and protozoa (3). There are three pathways of complement activation : classical, lectin, and alternative, all of which converge at the cleavage of complement component 3 (C3) (4). The alternative pathway distinguishes itself from the other two pathways by possessing a unique feature: a potent positive feedback loop that amplifies the activation of C3 regardless of the initial pathway involved (5). This amplification mechanism leads to an increased production of various pro-inflammatory effectors associated with complement (5). Notably, the alternative pathway is considered a driving force behind pathological complement activation, which contributes to the development of disease (5). A key factor in the regulation of the alternative pathway activation is properdin (FP), a positive regulator that facilitates and stabilises the assembly of C3bBb, the alternative pathway C3 convertase (6). Given that the alternative pathway is constitutively active at baseline and can also be activated because of classical/lectin pathway activation through the amplification loop, its regulation is crucial. In this regard, factor H (FH) plays a pivotal role in controlling the alternative pathway activity (7).

Previous studies on SARS-CoV have revealed its interaction with mannan-binding lectin (MBL), leading to the activation of the lectin pathway (8). Similarly, involvement of the complement system was shown in inducing hyperinflammation in middle east respiratory syndrome coronavirus (MERS-CoV) infection in animal models (9). Recently, attenuation of SARS-CoV-2 Infection has been shown by C1q and C4BP independently of complement activation (10).

SARS-CoV-2 has demonstrated the ability to activate the complement system through all three pathways (11). Dysregulation of the alternative pathway has been implicated in the severity and mortality associated with SARS-CoV-2 infection (12). Importantly, the alternative pathway can be directly activated by the SARS-CoV-2 spike protein (13). Patients with severe COVID-19 exhibit abnormality in gene expression levels of key alternative pathway proteins, including factor H (FH) and properdin (FP) (12, 14–16). Complement FH is a soluble glycoprotein with a molecular weight of 155 kDa and functions as a negative regulator in the alternative pathway (17, 18). FH is present in human plasma at concentrations ranging from 128 to 654 µg/ml (17, 18). FH is synthesised by various immune and non-immune cells, including epithelial and endothelial cells (18, 19). A previous study has demonstrated interactions between FH and the soluble West Nile virus NS1 protein, suggesting a potential role for FH in the immune response against West Nile virus infection (20). Additionally, FH has been found to bind and inhibit influenza A virus (IAV) cell entry and attenuate inflammatory responses in A549 cells in a subtype-dependent manner (6). These findings highlight the importance of FH in the immune defence against viral infections.

Human FP is critical in stabilising C3 convertases by forming complexes such as C3bBbP and C3bBbC3bP (4). This stabilisation prolongs the half-life of these complexes and enhances the alternative pathway activity (4). Unlike most complement proteins that are primarily synthesised in the liver, FP is produced at a range of locations by immune active cells (4). Neutrophils are a major source of FP secretion but FP is also synthesised by monocytes and T cells (7). In human serum, FP exists as cyclic polymers, including cyclic dimers, trimers, and tetramers, with a plasma concentration ranging from 22 to 25 µg/ml (7). The monomeric form of FP consists of seven non-identical thrombospondin type 1 repeats (TSRs) and has a molecular weight of 53 kDa. Among these TSRs, TSR4 and TSR5 are believed to be crucial for binding to C3bBb (7). In recent studies, a recombinant form of TSR4+5, produced as a double domain, has been shown to bind C3b and inhibit the alternative pathway activity (7). Moreover, FP has been found to inhibit influenza A virus (H1N1 subtype) cell entry and binding and attenuate the pro-inflammatory immune response in a complement activation - independent manner (7).

To date, the immune functions of FH and FP have not been explored in the context of SARS-CoV-2 infection, independent of their involvement in complement activation/modulation. This study employed purified human FH, FP and recombinant TSR4+5 proteins to investigate their potential protective or pathogenic roles in SARS-CoV-2 infection. We examined the interaction of FH and FP with SARS-CoV-2 S and RBD proteins. We assessed the potential of FH and FP to inhibit viral entry using SARS-CoV-2 pseudotype viral particles in A549 cells expressing human ACE2 and TMPRSS2 receptors/co-receptors. Our findings demonstrate that FH can effectively inhibit SARS-CoV-2 pseudotype cell entry, while FP enhances it. Moreover, we observed that treatment with FH resulted in the downregulation of the proinflammatory response triggered by SARS-CoV-2, whereas FP upregulated it.

## Materials and Methods

### Purification of human complement factor H (FH)

FH was isolated from human plasma using an affinity column made up of a monoclonal antibody against human FH (MRCOX23; MRC Immunochemistry Unit, University of Oxford) coupled to CNBr-activated Sepharose (GE Healthcare, UK) as described previously (21). Freshly thawed human plasma (50ml) (TCS Biosciences) was adjusted to a concentration of 5mM EDTA, pH 8 and dialysed against buffer I (25 mM Tris-HCL, 140 mM NaCl, and 0.5 mM EDTA at pH 7.5) with stirring overnight at 4°C. Dialysed plasma was passed through the MRCOX23 Sepharose column after it had been washed with 5 bed volumes of distilled water and buffer I. FH was then eluted using 3M magnesium chloride (Merck), 25 mM Tris-HCl, pH 8.0, 140 mM NaCl, and 0.5 mM EDTA. The collected fractions (1 ml each) were neutralised using 1 M Tris pH 7.5. The fractions were then dialysed against 1 L of distilled water overnight. The next day, the fractions were dialysed against H_2_O overnight, followed by 10 mM potassium phosphate for 4h. Western blotting was used to analyse the samples to determine their identity (Supplementary Figure S1A).

### Purification of native human properdin (FP)

Human native properdin was purified using a monoclonal antibody affinity column, as previously described (22, 23). Briefly, 100 ml plasma was made 5 mM EDTA, pH 8, centrifuged at 5000 x g for 10 min, and filtered using Whatman filter paper to remove debris/precipitate. An IgG-Sepharose column was equilibrated with 3-bed volumes of HEPES buffer (10 mM HEPES, 140 mM NaCl, 0.5 mM EDTA, pH 7.4). The plasma was then passed through the IgG-Sepharose column to remove C1q. The C1q-depleted plasma was passed through the anti-properdin antibody column and washed with 3 bed volumes of HEPES buffer. Next, the bound properdin was eluted using 3M MgCl_2_. The eluted fractions (1 ml each) were dialysed against HEPES buffer, stirring overnight at 4°C. Minor contaminants were removed by passing the fractions through the HiTrap Q FF-Sepharose ion exchange column (GE Healthcare, UK) to remove any impurities. Western blotting was used to evaluate immunoreactivity of the samples (Supplementary Figure S1B).

### Expression and purification of properdin-TSR4+5

Recombinant TSR4+5 nodule was expressed and produced in *E. coli* BL21 cells as a recombinant protein, fused to maltose-binding protein (MBP), using an amylose resin column, as previously described (22, 24). Briefly, 12.5 mL of the primary culture containing the protein-expressing cells was added to a 500 mL LB medium supplemented with 100 µg/mL of ampicillin. The culture was placed on a shaker and incubated at 37°C for 3h until the OD600 reached 0.6. After centrifugation at 13,800 x g for 10 minutes, the bacterial cell pellet was resuspended in 25 mL of a lysis buffer containing 20 mM Tris-HCl (pH 8.0), 0.5 M NaCl, 1 mM EGTA (pH 7.5), 1 mM EDTA (pH 7.5), 5% v/v glycerol, 0.2% v/v Tween 20, 0.1 mM PMSF, and 50 µg/mL lysozyme. The resuspended cells were incubated at 4°C for 30 minutes. Following incubation, the lysate was subjected to sonication with 12 cycles at 60 Hz for 30 seconds each, with a 2-minute interval between cycles. The lysate was then centrifuged at 15,000 x g for 30 min at 4°C. The supernatant obtained after centrifugation was mixed with 125 mL of buffer I, which contained 20 mM Tris-HCl (pH 8.0), 100 mM NaCl, 1 mM EDTA (pH 7.5), 0.2% v/v Tween 20, and 5% v/v glycerol. The diluted supernatant was loaded onto a 5 mL amylose resin column (New England Biolabs). The column was washed with 150 mL of buffer I, followed by 250 mL of buffer II (buffer I without Tween 20). The fusion protein was eluted using 100 mL of buffer II containing 100 mM maltose. The protein concentration was determined by measuring A_280_. The peak fractions were passed through Pierce™ High-Capacity Endotoxin Removal Resin (Qiagen, Hilden, Germany) to remove any remaining lipopolysaccharides (LPS). The levels of endotoxin in the purified protein samples were assessed using the QCL-1000 Limulus amebocyte lysate system (Lonza, Basel, Switzerland), and it was found that the recombinant proteins had endotoxin levels of ∼ 4 pg/µg. The purified proteins were quantified (yield ∼2 mg) and analysed using western blotting (Supplementary Figure S1C).

### Cell culture

Adenocarcinoma human alveolar basal epithelial cells (A549) expressing co-receptors, human ACE2 and TMPRSS2, used were produced, as previously described (10). Briefly, A549 cells were cultured in Dulbecco’s Modified Eagle’s Medium (DMEM) with Glutamax (Gibco) supplemented with 100U/ml penicillin (Gibco), 100µg/ml streptomycin, and 10% v/v foetal bovine serum (FBS) (Gibco) (complete growth media) at 37°C with 5% v/v CO_2_. Once the cells reached 70% confluence, they were transiently co-transfected with a plasmid expressing human ACE2 (pCDNA3.1+-ACE2) and another expressing TMPRSS2 (pCAGGS-TMPRSS2), using FuGENE™ HD Transfection Reagent (Promega). The following day, A549 cells were cultured in the presence of hygromycin and puromycin (Thermo Fisher Scientific, Waltham, MA, USA) to select the cells that co-expressed human ACE2 and TMPRSS2 (A549-hACE2+TMPRSS2 cells) on the cell surface.

### ELISA

To assess the binding of SARS-CoV-2 S or RBD proteins to either immobilised FH or FP, decreasing concentrations (1, 0.5, 1.25, and 0 µg/well) of immobilised FH or FP were coated on polystyrene microtiter plates (Sigma-Aldrich) overnight at 4°C using carbonate/bicarbonate (CBC) buffer, pH 9.6. As a negative control, wells were coated with 1µg/well BSA. Next day, the wells were washed 3 times with PBST Buffer (PBS + 0.05% Tween 20) (Fisher Scientific) to remove unbound proteins. Following the washes, 2% w/v BSA in PBS (Fisher Scientific) was used to block the wells for 2h at 37°C. The plate was then washed 3 times with PBST again to remove any excess BSA.

A constant dose of 1µg/well in 100 µl, recombinant SARS-CoV-2 S protein (RP-87680, Invitrogen) or recombinant SARS-CoV-2 spike RBD protein (40592-V08H, Sino-Biological) was added into the corresponding wells in which FH or FP was coated. Similarly, another experiment was performed to examine the binding of the FH or FP to the immobilised SARS-CoV-2 S or RDB. Decreasing concentrations of FH or FP (1, 0.5, 1.25, and 0 µg/well) were added to immobilised SARS-CoV-2 S (1 µg/100 µl) or RBD (1 µg/100 µl) coated wells. The wells were washed 3 times with PBST, then blocked with 2% w/v BSA in PBS for 2h at 37°C, and washed 3x with PBST.

The binding of the immobilised FH or FP were detected using polyclonal rabbit anti-SARS CoV-2 S (NR-52947, Bei-Resources), whereas, for the immobilised SARS-CoV-2 spike or RBD protein, the respective primary antibodies, rabbit anti-human properdin (MRC immunochemistry unit, Oxford) or monoclonal mouse anti-human factor H (MRCOX23) (MRC Immunochemistry Unit, Oxford) were used, each at a dilution 1:5000, and incubated for 1h at 37°C. The plates were washed 3 times using PBST to remove unbound antibodies. As secondary antibodies (1:5000), goat anti-rabbit IgG conjugated to horseradish peroxidase (HRP) (Promega) was added and incubated for one h at 37°C. Binding was detected by using a 3,3’,5,5’-tetramethylbenzidine (TMB) substrate set (Biolegend), and the reaction was stopped with 1M of H_2_SO_4_ (Sigma-Aldrich). iMark™ microplate absorbance reader (BioRad) was used to read the plate at 450nm.

### Viral cell entry assay

#### Treatment of SARS-CoV-2 Spike Protein Pseudotyped lentiviral particles

Pseudotyped lentiviral particles were produced as previously described (25). SARS-CoV-2 lentiviral pseudoparticles, pre-incubated with 20 µg/ml FH, FP (with or without 20 µg/ml anti-human FP polyclonal antibodies), or TSR4+5 for 2h at RT, were used to challenge A549-h ACE2+TMPRSS2 cells. Cells, challenged with lentiviral pseudoparticles without complement proteins, were used as untreated controls.

#### Luciferase Assay

An infection assay based on Luciferase reporter activity was used to examine whether FH, FP or TSR4+5 treatment may impact SARS-CoV-2 cell entry. Briefly, A549-hACE2+TMPRSS2 cells (20,000 cells/well) were seeded in a 96 well-plate and left to adhere overnight at 37°C in complete growth media. The next day, the cells were challenged with FH, FP or TSR4+5-treated SARS CoV-2 lentiviral pseudoparticles (as described above) in serum-free growth media (DMEM with Glutamax supplemented with 100U/ml penicillin, and 100µg/ml streptomycin) and incubated for 24h at 37°C. Next, the cells were washed 2 times with PBS, and fresh growth media was added and incubated for another 48h at 37°C. Post incubation, the cells were washed, and luciferase activity (RLU) was measured using the ONE-GloTM Luciferase Assay System (Promega) and Clariostar Plus Microplate Reader (BMG Labtech).

#### NF-κB activity assay

A luciferase-based reporter assay was used to assess the likely role of FH and FP in modulating NF-κB activity during SARS-CoV-2 infection. Briefly, A549-hACE2+TMPRSS2 cells were transfected with the pNF-κB-LUC plasmid (#631904, Clonetech). This vector comprises multiple copies of the NF-kB consensus sequences fused to a TATA-like promoter region of the Herpes Simplex Virus Thymidine Kinase (HSV-TK) promoter. The plasmid is engineered to measure the binding of the transcription factor to the κ enhancer providing a direct measurement of this pathway. After endogenous NF-κB binds to the κ-enhancer element, transcription is induced, and the reporter gene is activated. Using the Promega FuGENE^TM^ HD Transfection Reagent, the A549 cells were transfected with the plasmid and incubated for 48h at 37°C in complete growth media. Post transfection, the cells (20,000 cells/well) were plated in a 96 well-plate and left to adhere overnight at 37°C in complete growth media. The cells then were challenged with SARS-CoV-2 S protein (500ng/ml), pre-treated with 20 µg/ml FH or FP for 2h at RT. The cells were then incubated for 24h at 37°C in incomplete growth media. A549-hACE2+TMPRSS2 cells+SARS-CoV-2 S protein without complement protein pre-incubation was used as a control. Luciferase reporter assay for measuring NF-κB activity was carried out, as described above.

#### Cell binding assay

A cell binding assay was set up to assess whether FH or FP treatment affected SARS-CoV-2 cell binding. Briefly, A549-hACE2+TMPRSS2 cells (20,000 cells/well) were seeded in a 96 well-plate and left overnight at 37°C in growth media to adhere. The cells were then challenged with SARS-CoV-2 lentiviral pseudoparticles pre-treated with FH, FP (with or without 20 µg/ml anti-human FP polyclonal antibodies), or TSR4+5 (as described above), and incubated for 2h at 37°C in incomplete growth media. This was followed by washing the wells 3 times with PBS, then fixed for 1 min with 1% v/v paraformaldehyde (PFA) at RT. That was followed by 3 round of washing step with PBS and incubation with polyclonal rabbit anti-SARS-CoV-2 S (1:200) for 1h at 37°C. After washing again 3 times with PBST, the respective wells were probed with Alexa Fluor 488 conjugated goat anti-rabbit antibody (1:200) (Abcam) for 1h at RT. Clariostar Plus Microplate Reader (BMG Labtech) was used to read the plate.

### Estimating FH/FP modulated cytokine response during SARS-CoV-2 infection

#### SARS-CoV-2 alphaviral pseudoparticles treatment

SARS-CoV-2 alphaviral pseudoparticles that encoded the four structural proteins, S, E, M, and N (Ha-CoV-2 Luc; Virongy, Manassas, VA, USA), were utilised for gene expression analysis (RT-qPCR). Alphaviral pseudoparticles were pre-incubated with 20 µg/ml FH, FP or TSR4+5 for 2h at RT, and were then challenged against A549-hACE2+TMPRSS2 cells. Cells challenged with pseudoparticles without prior incubation with complement proteins were used as the untreated control.

#### Quantitative qRT-PCR analysis

qRT-PCR assay was performed to evaluate if FH or FP treatment may affect pro-inflammatory gene expression levels in cells challenged with SARS-CoV-2 pseudoparticles. Briefly, A549-hACE2+TMPRSS2 cells (0.5 × 10^6^) were seeded in a 12-well plate and left to adhere overnight at 37°C and 5% v/v CO_2_ in complete growth media. Next day, SARS-CoV-2-alphaviral pseudoparticles, pre-treated with FH, FP or TSR4+5 (as described above), were added to A549-h ACE2 + TMPRSS2 cells and incubated for 6 and 12h at 37°C in incomplete growth medium. The cells were then washed with PBS before being pelleted. GenElute Mammalian Total RNA Purification Kit (Sigma-Aldrich) was used to extract the total RNA, as per the manufacturer’s instructions. A Nanodrop 2000/2000c (ThermoFisher) was utilised to determine the amount of RNA at 260 nm after DNA contaminants were eliminated using DNase I (Sigma-Aldrich). The purity of the RNA was assessed using A260/A280 ratio. 2 μg of total RNA was used to synthesise cDNA using the High-Capacity RNA to cDNA Kit (Applied Biosystems). The primer sequences were generated using the primer BLAST software (Basic Local Alignment Search Tool)

Table *1*. The qRT-PCR assay was performed using Step One Plus equipment (Applied Biosciences). Each qPCR experiment in triplicates used 500 ng of cDNA, 75 nM forward and reverse primers, and 5 µl of Power SYBR Green Master Mix (Applied Biosystems). qPCR samples were run at 50°C and 95°C for 2 and 10 min, followed by the amplification template for 40 cycles, each cycle involving 15 s at 95°C and 1 min at 60°C. 18S rRNA was used as an endogenous control to normalise gene expression.

**Table 1:**
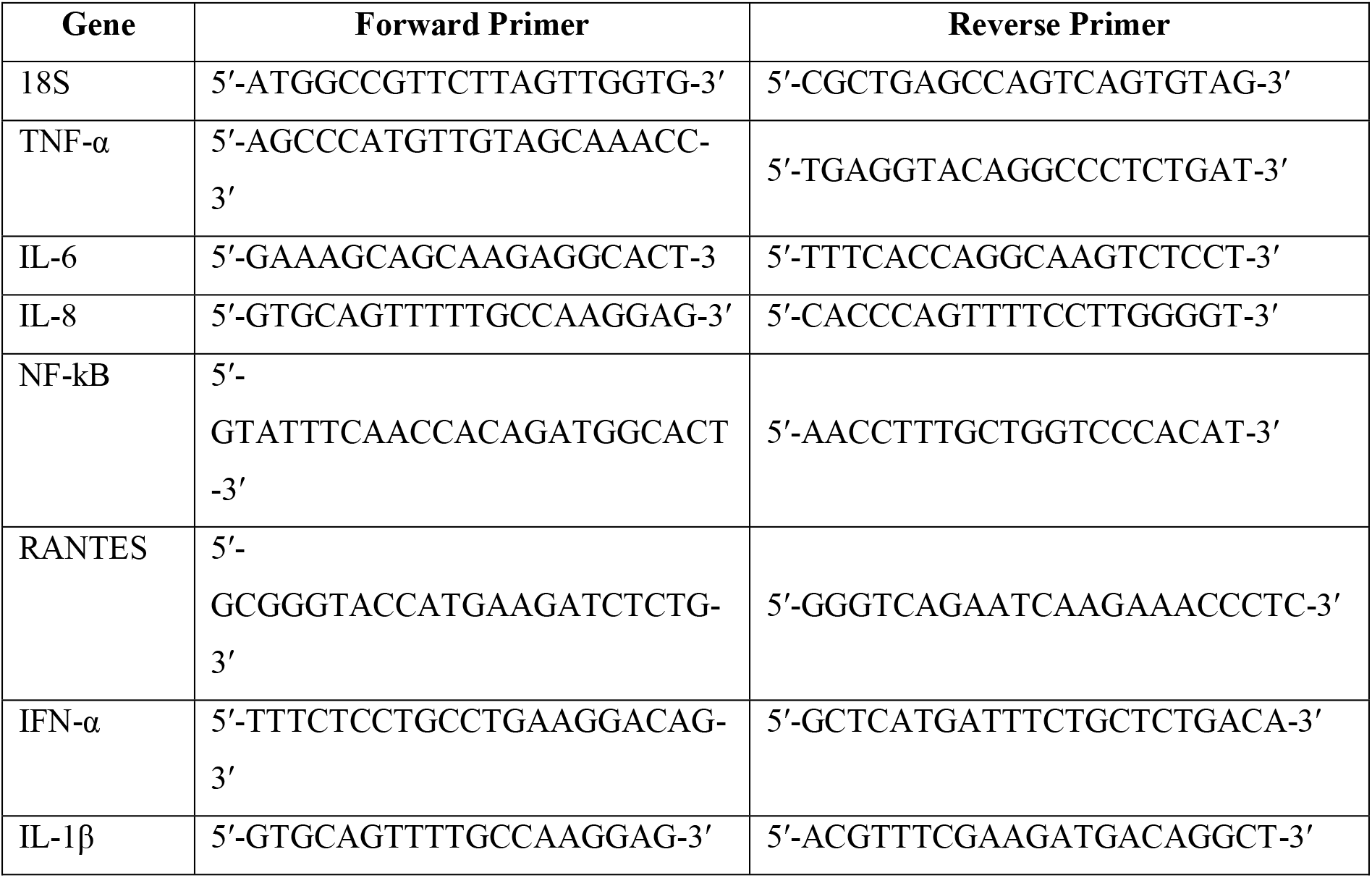
Forward and reverse primers used for qRT-PCR assay.

#### In silico interaction analysis involving FP, spike and ACE2 receptor

The binding site prediction in S protein for FP was performed using a blind docking approach. Structural coordinates for FP monomer (domains 1, 4, 5 & 6) and spike trimer were retrieved from RCSB with PDB IDs 6S08 and 6XM3, respectively. The electron microscopy structure of spike bound to ACE2 receptor (PDB ID: 7KNB) was retrieved and the inter-molecular interactions were analysed. The interacting residues were used to define the binding site for re-docking experiment with the aim to generate the binding energy of the docked complex. The previously docked complex of spike and FP, along with the ACE2 receptor structure, extracted from PDB ID: 7KNB, was used to generate a tripartite complex structure of spike, FP and ACE2. All proteins were prepared for docking using the ‘Prepare Proteins’ module of Discovery Studio (DS) 2021 with default parameters setting. ‘ZDOCK’ module of DS 2021 was used for docking with default parameters and docked poses were further refined using ‘ZRank’ algorithm. Top cluster poses were analysed, and the final pose was selected based on concurrence with known interaction of ACE2 with spike (as present in PDB ID: 7KNB), and maximal interaction of domains 4 (TSR4) and 5 (TSR5) of FP with spike. Binding free energy for three complexes, (a) spike and FP, (b) spike and ACE2, and (c) spike-FP and ACE2 were calculated using in-house ‘Binding Free Energies’ protocol in DS 2021.

An attempt was made to predict the intermolecular interactions of FH, spike, and ACE2 receptor. This experiment could not be performed due to unavailability of full-length 3D structure of FH in PDB. The Alpha-fold (ID: AF-P08603) or homology based modelled structure that was generated for FH was sub-optimal due to several unfolded regions and thus, was not fit for the docking studies (Supplementary Figure S3).

#### Statistical Analysis

The graphs were made with the aid of GraphPad Prism 9.0. According to the figure legends, the statistical significance between the treated and untreated situations was considered. The SD or SEM that the error bars reflect is stated in the figure legends.

## Results

### SARS-CoV-2 spike and RBD proteins bind to human FH and FP

A direct ELISA assay was employed to investigate the binding interactions between purified FH (Supplementary Figure S1A) and SARS-CoV-2 S and RBD proteins (Figure 1A), as well as the reciprocal binding of SARS-CoV-2 S and RBD proteins to immobilised FH Figure 1B). The results revealed a dose-dependent binding of FH to S and RBD proteins when probed with the anti-SARS-CoV-2 S protein polyclonal antibody. Similarly, immobilised S protein or RBD displayed a dose-dependent binding to FH when probed with the rabbit anti-human FH polyclonal antibody.

**Figure 1.**
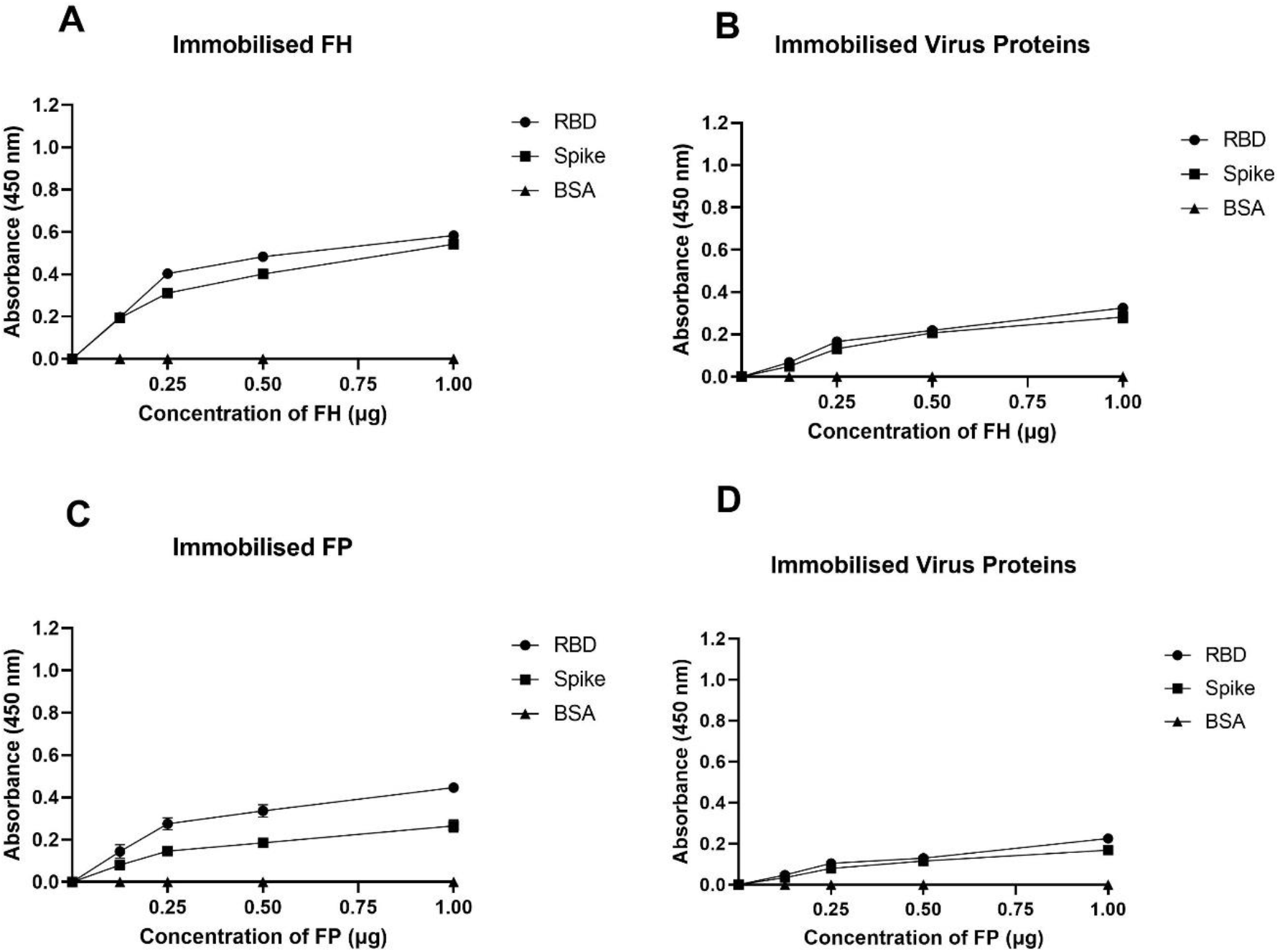
SARS-CoV-2 Spike protein directly interacts with FH and FP via its RBD. FH bound both SARS-CoV-2 Spike and RBD proteins in a dose-dependent manner. Decreasing concentration of FH or FP (1, 0.5, 1.25, and 0 μg/well) (A) and (C), or constant concentration of viral proteins (1 μg/well) (B) and (D) were immobilised using a 96-well plate using Carbonate-Bicarbonate (CBC) buffer, pH 9.6 at 4°C overnight. After washing off the excess CBC buffer with PBS, a constant concentration of viral proteins (1 μg/well) (A) and (C), or decreasing concentrations of FH or FP (1, 0.5, 1.25, and 0 μg/well) (B) and (D) were added to the corresponding wells, followed by incubation at 37°C for 2h. After washing off the unbound proteins, the wells were probed with corresponding primary antibodies (1:5000; 100 μl/well), i.e. rabbit anti-SARS-CoV-2 Spike or rabbit anti-human FH or FP polyclonal antibodies. BSA was used as a negative control. The data are presented as a mean of three independent experiments carried out in triplicates ±SEM.

The binding capacity of purified FP (Supplementary Figure S1B) to SARS-CoV-2 S and RBD proteins was assessed using the direct ELISA assay (Figure 1C), along with the reciprocal binding of SARS-CoV-2 S and RBD proteins to immobilised FP (Figure 1D). The results demonstrated that immobilised FP exhibited dose-dependent binding to S and RBD proteins when probed with the polyclonal anti-SARS-CoV-2 S protein antibody. Similarly, the immobilised SARS-CoV-2 S or RBD proteins showed dose-dependent binding to FP when probed with the rabbit anti-human FP polyclonal antibody. BSA was used as a negative control protein in the assay. A comparable result was obtained using recombinant TSR4+5 modules (Supplementary Figure S1C) when probed with anti-MBP antibodies (Supplementary Figure S2).

### FH restricted SARS-CoV-2 Pseudoparticle transduction while FP and TSR4+5 promoted

A luciferase reporter assay was conducted to assess the impact of FH, FP and TSR4+5 on SARS-CoV-2 infectivity. SARS-CoV-2 lentiviral pseudoparticles, treated with FH showed reduced viral transduction in A549-hACE2+TMPRSS2 cells while increased transduction was observed in the case of FP or TSR4+5 compared to their respective controls. Specifically, A549-hACE2+TMPRSS2 cells challenged with SARS-CoV-2 lentiviral pseudoparticles pre-treated with FH, significantly reduced viral infection by ∼ 25% (Figure 2A). FP and TSR4+5 treatment increased viral entry by ∼80% and ∼140% (Figures 2B, 2C), respectively, compared to their respective controls. These findings suggested that FH acted as an entry inhibitor for SARS-CoV-2 pseudotyped particles, whereas FP facilitated viral entry.

**Figure 2.**
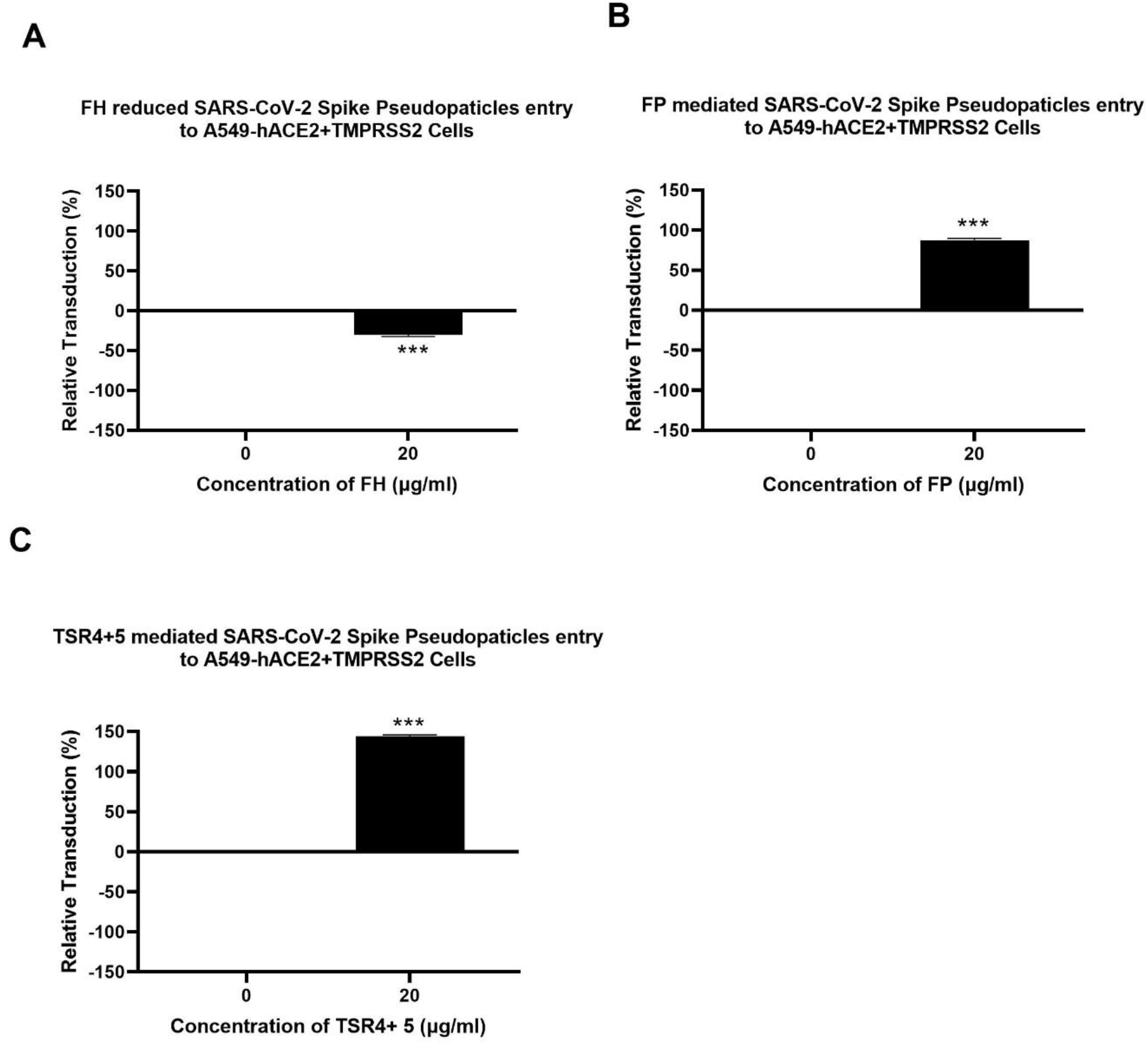
Modulation of SARS-CoV-2 pseudotype viral entry in A549-hACE2+TMPRSS2 cells by FH, FP or TSR4+5 treatment. FH, FP or TSR4+5 (20μg/ml) were used to pre-treat SARS CoV-2-lentiviral pseudoparticles (A), (B) and (C), respectively. To determine if the treatment affected the virus capacity to enter the cells, either treated or untreated lentiviral pseudoparticles were transduced into A549-hACE2+TMPRSS2 cells and then examined for luciferase reporter activity. The background was subtracted from all data points. The data obtained were normalised with 0% luciferase activity defined as the mean of the relative luminescence units recorded from the control sample (Cells + lentiviral pseudoparticles). Data are shown as the normalised mean of three independent experiments done in triplicates ±SEM. Significance was determined using the two-way ANOVA test (***p < 0.001) (n = 3).

### SARS-CoV-2 Pseudoparticle binding to the target cells was inhibited by FH and enhanced by FP and TSR4+5

A cell binding assay was conducted to evaluate the effect of FH, FP and TSR4+5 on SARS-CoV-2 binding to lung epithelial-like cells. A549-hACE2+TMPRSS2 cells were challenged with SARS-CoV-2 lentiviral pseudoparticles treated with FH, FP, and TSR4+5. Pre-treatment of SARS-CoV-2 pseudoparticles with FH decreased viral binding by ∼35% (Figure 3A). In contrast, compared to the control, FP and TSR4+5 promoted the binding by∼ 30% and∼ 50%, (Figures 3B, 3C), respectively. Anti-FP antibody treatment mitigated the effect of FP on viral entry and binding by ∼ 98 % and ∼ 85 % (Figures 4A, 4B), respectively. These results imply that FH and FP modulate SARS-CoV-2 viral binding, viral entrance, and subsequent infection to lung epithelial-like cells in an antagonistic and complement activation-independent manner. Furthermore, sequestering or neutralising FP may limit viral binding and entry in SARS-CoV-2 infection, implicating the potential of anti-FP antibodies to limit the severity of the disease.

**Figure 3.**
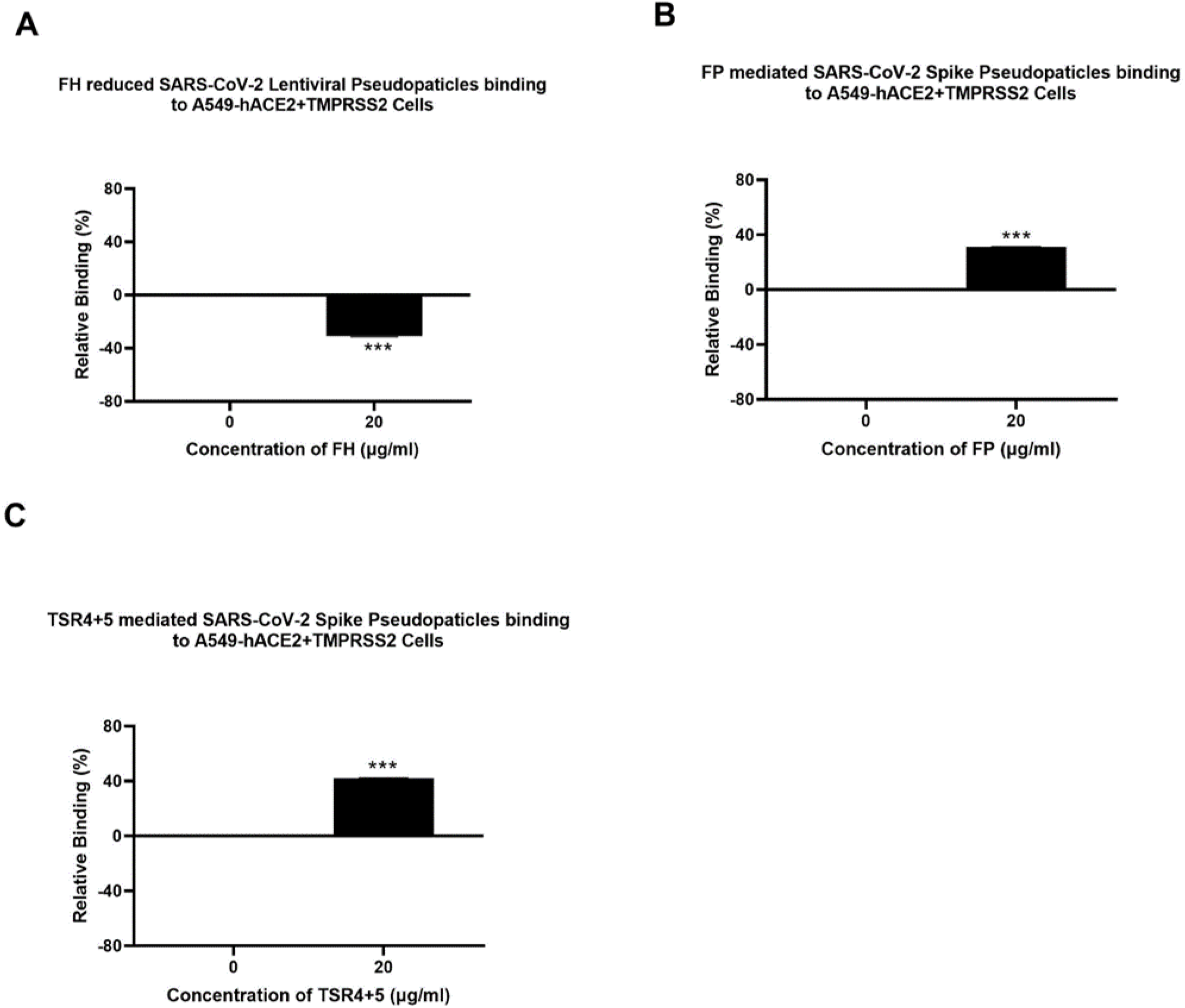
Modulation of SARS-CoV-2 pseudoparticle binding to A549-hACE2+TMPRSS2 cells by FH, FP or TSR4+5. Cell binding assay revealed FH treatment inhibits the binding to the cell-surface receptor, whereas FP and TSR4+5 mediated it (A), (B), and (C), respectively. A549-hACE2+TMPRSS2 cells (1 × 10^5^ cells/ml) were challenged with SARS-CoV-2 lentiviral pseudoparticles pre-incubation with or without FH, FP or TSR4+5 (20 μg/ml), followed by incubating at 37°C for 2h. After removing unbound protein and viral particles, the wells were fixed with 1% v/v paraformaldehyde for 1 min and probed with polyclonal rabbit anti-SARS CoV-2 spike (1:200). The data obtained were normalised with 0% fluorescence defined as the mean of the relative fluorescence units recorded from the control sample (Cells + lentiviral pseudoparticles). Three independent experiments were carried out in triplicates; error bars are expressed as ± SEM. Significance was determined using the two-way ANOVA test (***p < 0.001) (n = 3).

**Figure 4.**
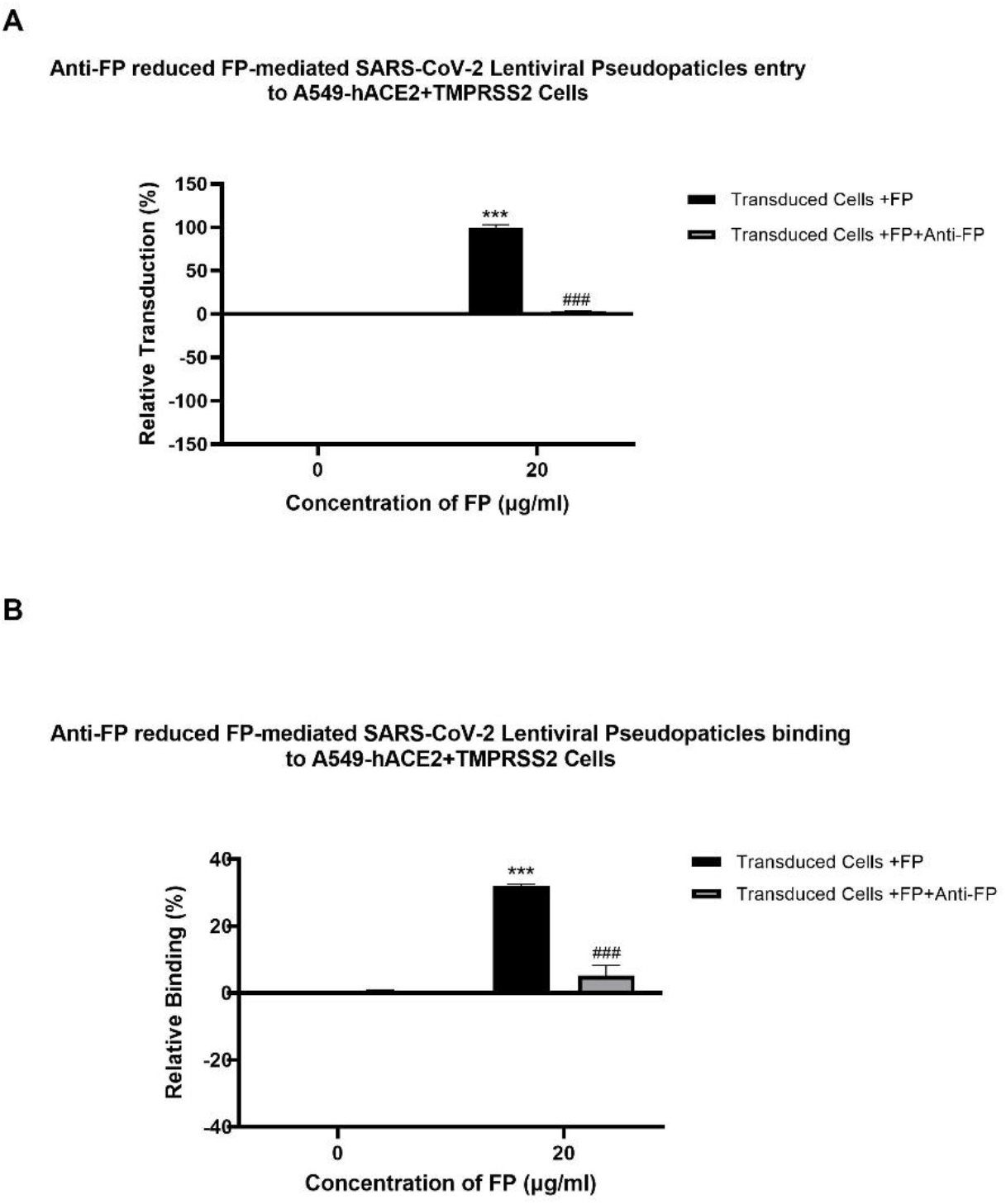
Reversal of FP mediated-SARS-CoV-2 viral entry in and binding to A549-hACE2+TMPRSS2 cells by anti-FP antibody. FP-treated (with or without anti-FP antibodies) or untreated lentiviral pseudoparticles were added to A549-hACE2+TMPRSS2 cells and luciferase reporter activity or cell binding were assessed. The background was subtracted from all data points. The data obtained were normalised with 0% luciferase activity defined as the mean of the relative luminescence units recorded from the control sample (Cells + lentiviral pseudoparticles). Data are shown as the normalised mean of three independent experiments done in triplicates ±SEM. Significance of FP-treated cells (with and without anti-FP antibodies) compared to the control (cells+ viral pseudoparticles) was determined using the two-way ANOVA test (***p < 0.001). Additionally, the significance of FP treated cells (with anti-FP antibodies) with respect to cells treated only with FP (cells+ viral pseudoparticles + FP) (p < 0.001) (n = 3).

### SARS-CoV-2 infection-associated inflammation can be attenuated by FH but promoted by FP

Since the NF-κB pathway is often associated with pro-inflammatory signals and responses, the effect of FP or FH treatment on NF-κB activation in lung epithelial-like cells challenged with SARS-CoV-2 was evaluated using NF-κB luciferase reporter assay. FP pre-treated SARS-CoV-2 spike protein exhibited a ∼60% increase in NF-κB activation in A549-hACE2+ TMPRSS2 cells, while FH showed ∼ 25% reduction in NF-κB activation compared to the control (Figures 5A, 5B), respectively.

**Figure 5.**
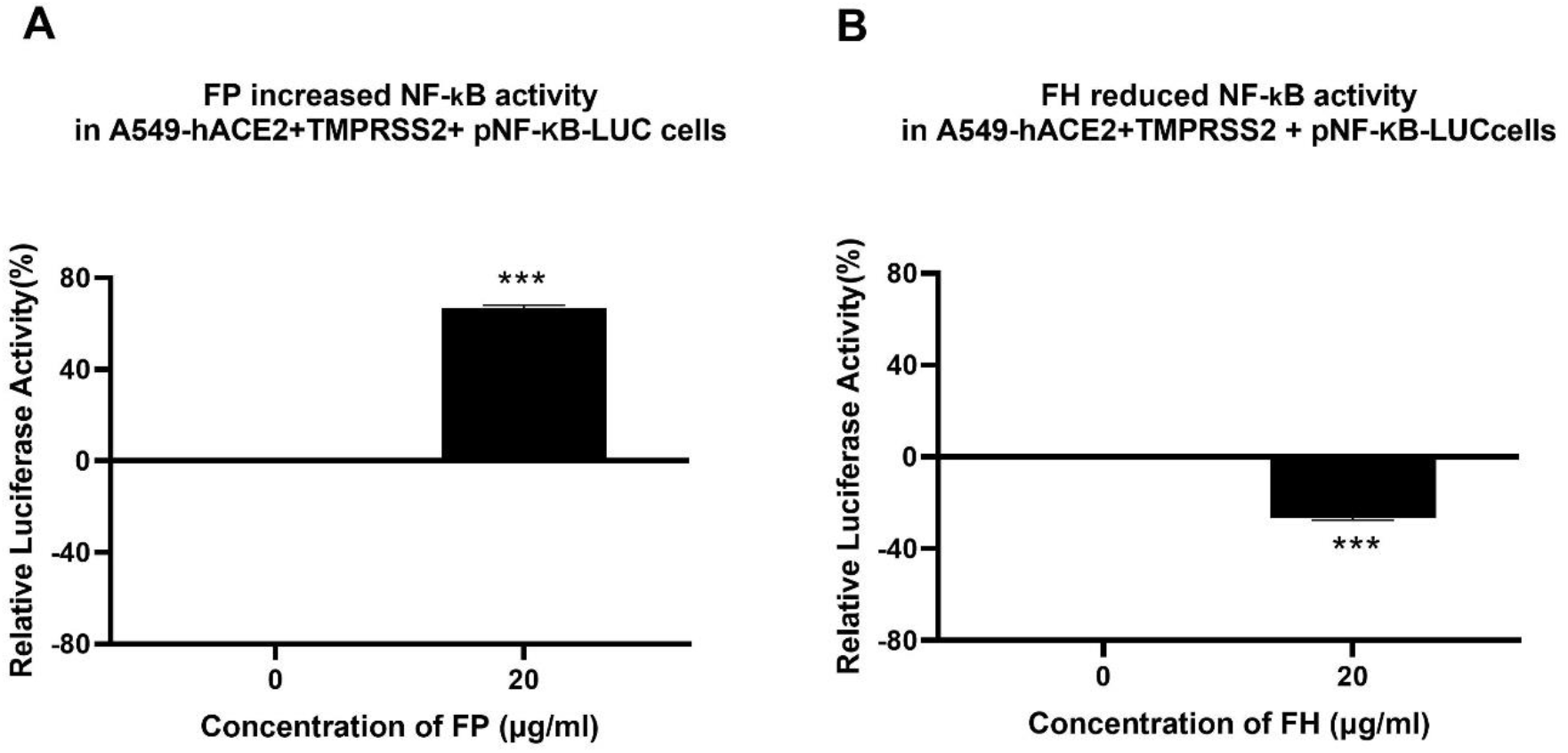
FP increases while FH reduces NF-κB activation in SARS-CoV-2-Spike protein challenged A549-hACE2-TMPRSS2 cells. SARS CoV-2-Spike protein pre-treatment with FP (A) or FH (B) altered NF-kB activation. To examine the immunological role of FP and FH on NF-kB activation, A549-hACE2+TMPRSS2 cells transfected with pNF-κB-LUC were challenged with SARS-CoV-2 Spike protein (500ng/ml) after pre-treatment with FP or FH (20μg/ml). The cells were then incubated for 24h and examined for luciferase reporter activity. The background was subtracted from all data points. The data obtained were normalised with 0% luciferase activity defined as the mean of the relative luminescence units recorded from the control sample (A549-hACE2+TMPRSS2 cells + SARS CoV-2-Spike protein). Data are demonstrated as the normalised mean of three independent experiments done in triplicates ± SEM. Significance was determined using the two-way ANOVA test (***p < 0.001) (n = 3).

Using qRT-PCR assay, we examined if FH, FP and TSR4+5 modulated cytokine response during SARS-CoV-2 infection. This was done by comparing the mRNA levels of pro-inflammatory cytokines and chemokines in lung epithelial-like cells challenged with SARS CoV-2-alphaviral pseudoparticles pre-treated FH, FP or TSR4+5, with their corresponding controls. The results showed that FH, FP and TSR4+5 modulated inflammatory immune response differentially in A549-hACE2+TMPRSS2 cells (Figures 6, 7 and 8), respectively. mRNA levels of pro-inflammatory cytokines such as IFN-α, IL-6, RANTES, IL-1β, IL-8 and TNF-α (and NF-κB) were downregulated in A549-hACE2+ TMPRSS2 cells challenged with SARS CoV-2 alphaviral pseudoparticles pre-treated with FH (FH-treated cells) compared to control cells (Figure 6).

**Figure 6.**
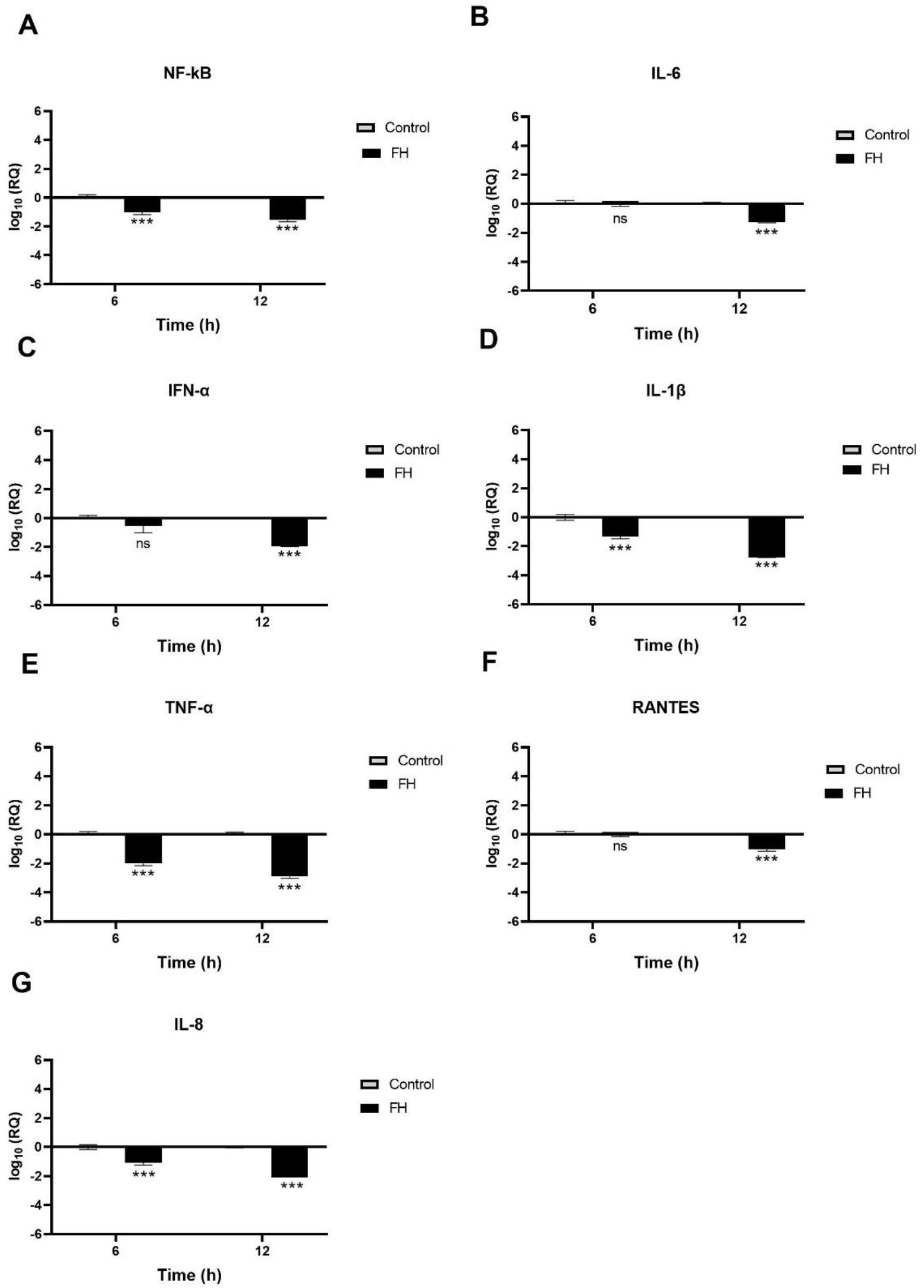
SARS-CoV-2 associated inflammation in A549-hACE2+TMPRSS2 cells is attenuated by FH. A549-hACE2+TMPRSS2 cells were challenged with and without FH (20μg/ml) pre-treated SARS-CoV-2 alphaviral pseudoparticles. Cytokines and chemokines mRNA levels were assessed using RT-qPCR for NF-κB (A), IL-6 (B), IFN-α (C), IL-1β (D), TNF-α (E), RANTES (F) and IL-8 (G). The relative expression (RQ) was calculated using the untreated cells (A549-hACE2+TMPRSS2 cells + SARS-CoV-2 alphaviral pseudoparticles) as the calibrator. The RQ value was calculated using the formula: RQ= 2^-ΔΔCt^. Experiments were carried out in triplicates, and error bars represent ± SEM (n =3); Significance was determined using the two-way ANOVA test (****p < 0.0001).

**Figure 7.**
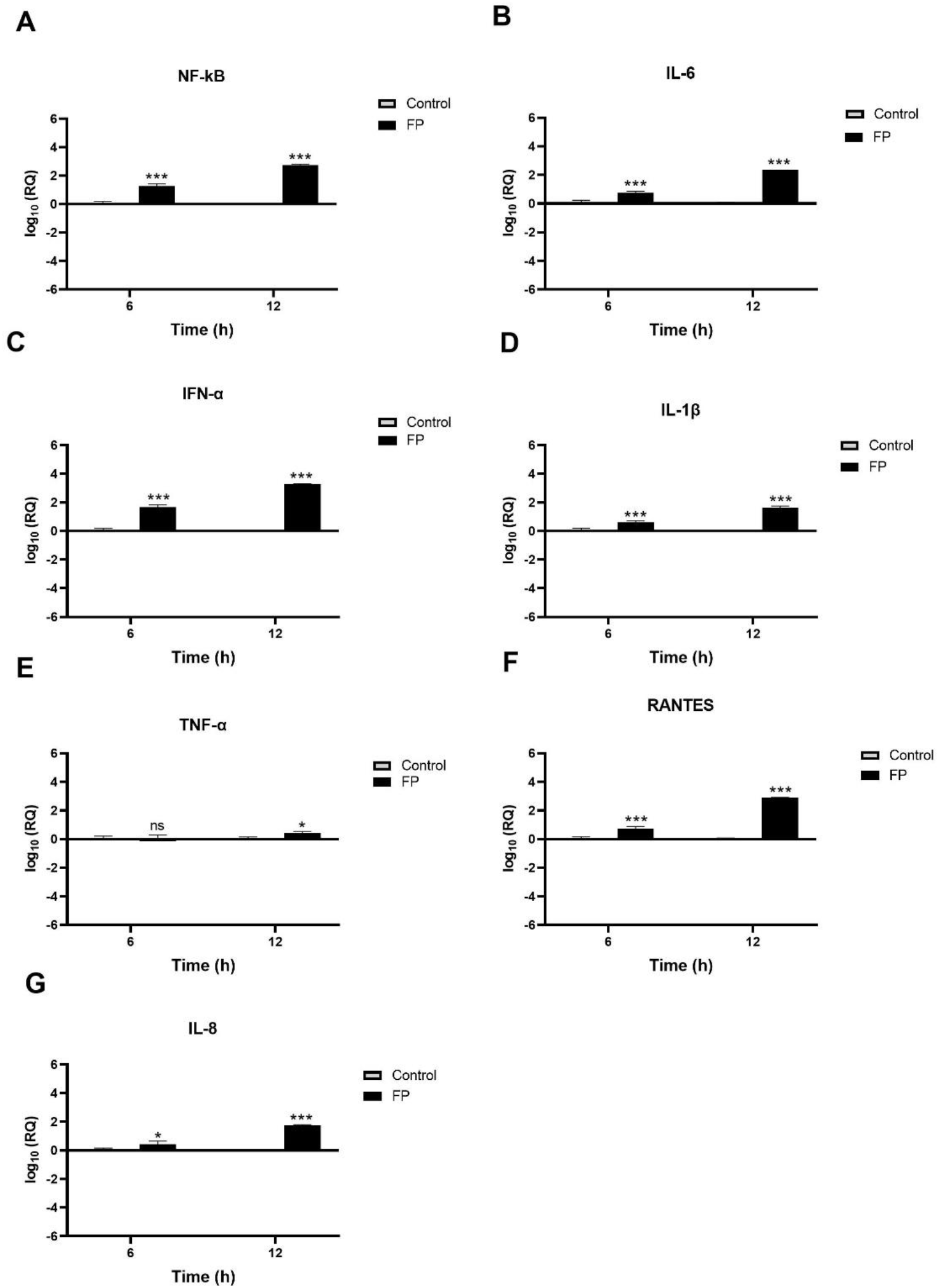
SARS-CoV-2 associated inflammation in A549-hACE2+TMPRSS2 cells is promoted by FP. FP pre-treated SARS-CoV-2 alphaviral pseudoparticles induces a pro-inflammatory response at 6h and 12h post infection in A549-hACE2 +TMPRSS2 cells. mRNA expression levels of targeted cytokines and chemokines of NF-κB (A), IL-6 (B), IFN-α (C), IL-1β (D), TNF-α, RANTES (F) and IL-8 (G) were measured using RT-qPCR. The data were normalised through 18S rRNA expression as an endogenous control. The relative expression (RQ) was calculated using the untreated cells (A549-hACE2+TMPRSS2 cells + SARS-CoV-2 alphaviral pseudoparticles) as the calibrator. The RQ value was calculated using the formula: RQ= 2^-ΔΔCt^. Assays were conducted in triplicates, and error bars represent ±SEM. Significance was determined using the two-way ANOVA test (*p 0.05, and ****p < 0.0001) (n = 3).

**Figure 8.**
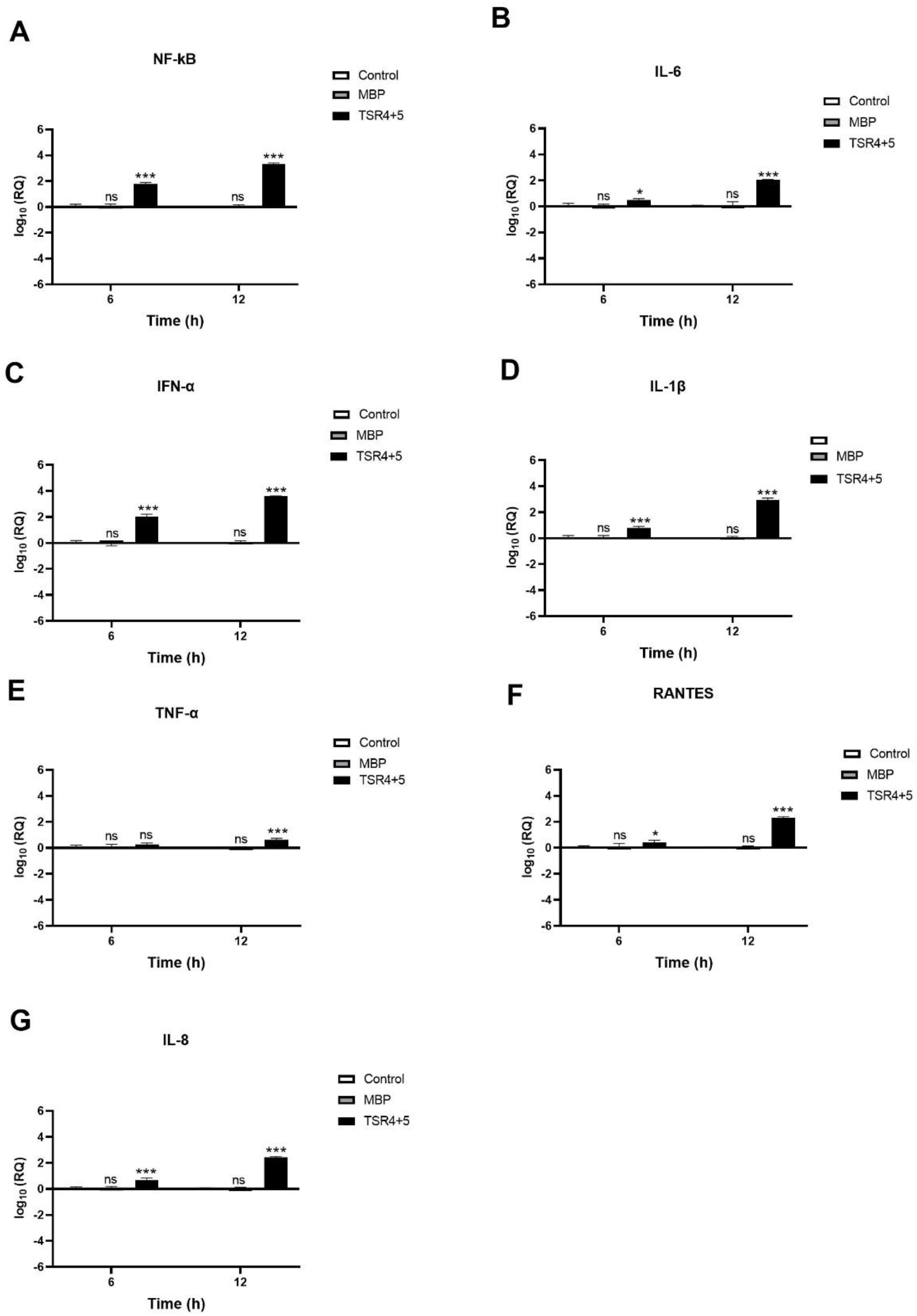
SARS-CoV-2 infection of A549-hACE2+TMPRSS2 cells induced greater pro-inflammatory response in the presence of TSR4+5. Pro-inflammatory events were triggered in response to TSR4+5 pre-treated SARS-CoV-2 alphaviral pseudoparticles in A549-hACE2 + TMPRSS2 cells at 6h and 12h post infection. The levels of gene expression of the cytokine and chemokines production were measured using qRT-PCR for NF-kB (A), IL-6 (B), IFN-α (C), IL-1β (D), TNF-α (E), RANTES (F) and IL-8 (G). The data were normalised against 18S rRNA expression as a control. Experiments were conducted in triplicates, and error bars represent ±SEM. The relative expression (RQ) was calculated using A549-hACE2 + TMPRSS2 cells + SARS-CoV-2 alphaviral pseudoparticles untreated with TSR4+5 as the calibrator. RQ = 2^-ΔΔCt^ was used to calculate the RQ value. Significance was determined using the two-way ANOVA test (*p 0.05, and ****p < 0.0001) (n = 3).

NF-κB gene expression levels decreased in FH-treated cells at 6 h (∼ −0.9 log10), peaking at 12h (∼ −1.7 log10) (Figure 6A). FH-treated cells at 6h exhibited a reduction in the gene expression levels of IL-1β (∼ −1.3 log10) (Figure 6D), TNF-α (∼ −2.0 log10) (Figure 6E), and IL-8 (∼ −0.1 log10) (Figure 6G) compared to their respective controls. Similarly, at 12h post-infection, FH-treated cells showed decreased mRNA levels of IL-6 (∼ −1.2 log10) (Figure 6B), IFN-α (∼ −1.9 log10 (Figure 6C)), IL-1β (∼ −2.7 log10) (Figure 6D), and IFN-α (∼ −2.9 log10), RANTES (∼ −0.8 log10) (Figure 6F), and IL-8 (∼ −2.1 log10) (Figure 6G) compared to untreated cells (Figure 6E). There were no significant changes in the mRNA levels of IL-6 at 6 h (Figure 6B), IFN-α (Figure 6C), and RANTES (Figure 6F) in FH-treated cells as compared to their controls.

A549-hACE2+TMPRSS2 cells, challenged with SARS CoV-2 alphaviral pseudoparticles that were pre-treated with FP or TSR4+5, the pro-inflammatory immune response was found to be upregulated (Figures 7, 8). NF-κB gene expression level in FP-treated cells was increased (∼ 0.7 log10) at 6 h than in control cells, and it was even higher (∼2.8 log10) at 12 h (Figure 7A). At 6 h, IL-6 (∼0.7 log10) (Figure 6B), IFN-α (∼1.5 log10) (Figure 7C), IL-1β (∼0.5 log10) (Figure 7D), RANTES (∼0.7 log10) (Figure 7F), and IL-8 (∼0.3 log10) (Figure 7G) were found to be downregulated in FP-treated cells. At 12 h, the mRNA levels of IL-6 (Figure 7B), IFN-α (Figure 7C), IL-1β (Figure 7D), TNF-α (Figure 7D), RANTES (Figure 7F), and IL-8 (Figure 7G) in FP-treated cells, were even more upregulated (∼2.7 log10, ∼3.4 log10, ∼1.5 log10, ∼0.2 log10, ∼3.0 log10, and ∼1.4 log10, respectively). There was no significant alteration in the mRNA levels of IFN-α at 6h in FP-treated cells respective to the control (Figure 7E).

Similarly, TSR4+5 - treated cells showed elevated NF-κB gene expression levels (∼0.2 log10) than control cells, with even higher levels at 12 h (∼2.0 log10) (Figure 8A). There was no significant difference between the control and the MBP-treated cells (TSR4+5 fusion protein) (Figure 8). Compared to their respective controls, at 6 h, the mRNA levels of IL-6 (∼0.5 log10) (Figure 8B), IFN-α (∼1.9 log10) (Figure 8C), IL-1β (∼0.7 log10) (Figure 8D), TNF-α (∼0.1 log10) (Figure 8E), RANTES (∼0.2 log10) (Figure 8F), and IL-8 (∼0.8 log10) (Figure 8G) in TSR4+5-treated cells were augmented as compared to the control. The downregulation in mRNA levels of IL-6 (∼ 2.2 log10) (Figure 8B), IFN-α (∼3.7 log10) (Figure 8C), IL-1β (∼2.7 log10) (Figure 8D), TNF-α (∼0.6 log10) (Figure 8E), RANTES (∼2.3 log10) (Figure 8F), and IL-8 (∼2.2 log10), were considerably more significant at 12h in TSR4+5-treated cells as compared to the untreated control. These findings suggest that FP and FH differentially modulate NF-κB activation and associated inflammatory response in SARS-CoV-2 infection.

### FP interacts with spike and ACE2 in a tripartite complex

A blind docking approach was attempted to generate a complex of FP and spike protein. The second ranked docked pose concurred with the *in vitro* observation that TSR4 and 5 domains of FP interacted with RBD and NTD domains of spike protein, respectively through H-bonding, electrostatic and hydrophobic interactions (Table 2; Figure 9A). A tripartite complex structure of FP, spike and ACE2 was created by docking electron microscopy structure of ACE2 to FP bound spike protein. In the first ranked pose, ACE2 was found to interact with spike as reported in the electron microscopy structure (PDB ID: 7KNB). It was observed that FP interacted with both spike and ACE2 in the tripartite complex through various non-bonded contacts in each subunit (Table 2; Figure 9B, C). In this structure, TSR4 domain of FP was also found proximal to ACE2 receptor and showed interaction with both spike and ACE2. The binding affinity of ACE2 receptor with unbound and FP-bound spike proteins was compared through Zdock score and binding free energy. These scores indicated that ACE2 receptor had a strong affinity for FP-bound spike as compared to unbound spike protein (Table 3). This suggests that FP may enhance the affinity of spike for ACE2 by interacting with both proteins through a tripartite complex.

**Figure 9.**
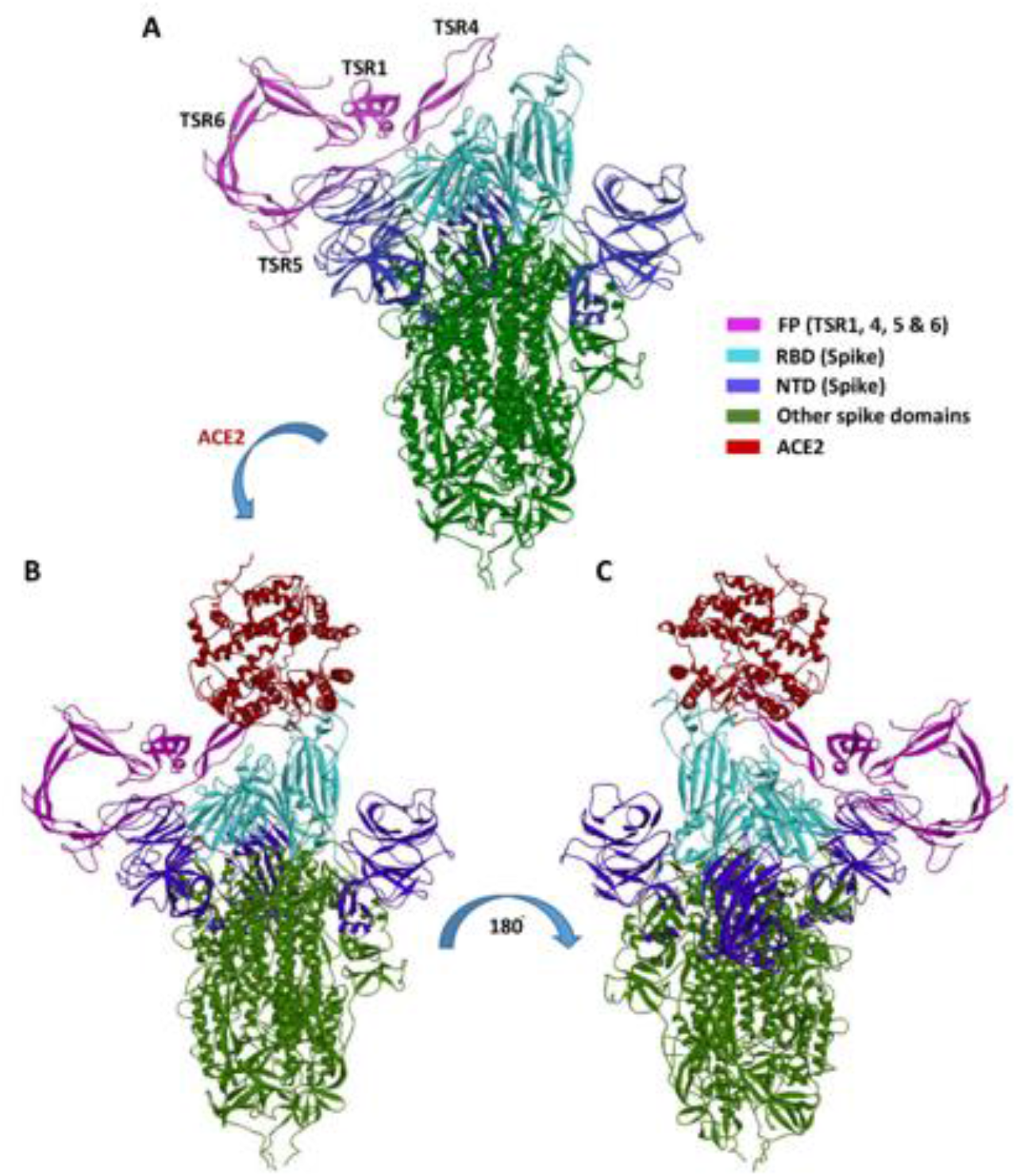
Cartoon representation of FP interaction with spike and ACE2. (A) Interaction of FP with spike RBD and NTD through TSR4 and TSR5 domains. (B) & (C) A tripartite complex representation of FP, spike and ACE2.

**Figure 10:**
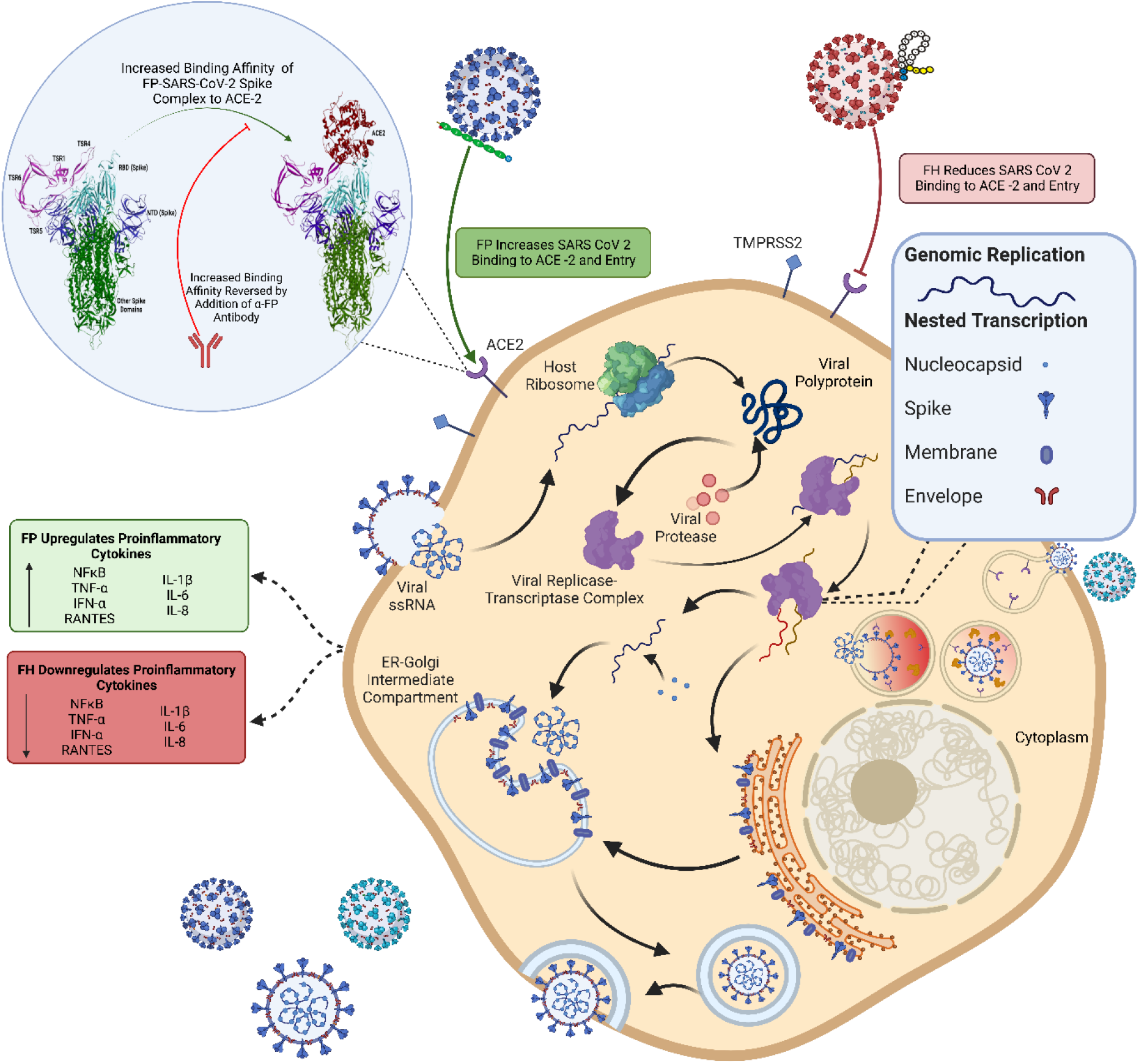
Immune Effector Function of Factor P (FP) and Factor H (FH) in SARS-CoV-2 Infection Independent of Complement Activation. The SARS-CoV-2 virus binds to host cells through the ACE2 receptor using its spike protein. After fusion with the cell membrane, the viral RNA is released into the host cytoplasm, initiating viral replication, protein synthesis and viral release into the extracellular environment contributing to the spread of SARS-CoV-2 infection. In this study, FP and FH were investigated separately to understand their effects on SARS-CoV-2 infection without activating the complement cascade. FP was found to enhance the binding affinity between the SARS-CoV-2 spike protein and ACE2, leading to increased viral entry into host cells and subsequent infection. This increased viral load triggered the upregulation of proinflammatory cytokines, contributing to the inflammatory response. Conversely, FH was observed to reduce the binding and entry of SARS-CoV-2 into host cells. Consequently, the lower viral entry mediated by FH resulted in decreased production of proinflammatory cytokines, potentially mitigating the immune response. These findings shed light on the immunomodulatory role of FH and the immunopathological role of FP in COVID-19. Moreover, they provide insights into the potential association between elevated levels of properdin and insufficient levels of FH observed in severe COVID-19 patients. This figure was modelled based on a published figure with permission of the rights holder, Elsevier GmbH, from Varghese et al., 2020 [1]. Created with BioRender.com.

**Table 2:** Interaction details of docked complexes of FP, Spike and ACE2.

| Receptor | Ligand | H-bonding residues |  | Electrostatic interaction residues |  | Hydrophobic interaction residues |  |
| --- | --- | --- | --- | --- | --- | --- | --- |
|  |  | Receptor | Ligand | Receptor | Ligand | Receptor | Ligand |
| Spike | FP | ASN122,<br>ASN125,<br>TYR145,<br>THR345,<br>LYS356,<br>LYS444 | CYS269,<br>GLN281,<br>ASP356,<br>GLN363,<br>GLN364,<br>GLN365,<br>ASN428 | PHE157 | ASP366 | LYS129,<br>CYS131,<br>CYS166,<br>THR167,<br>PHE168,<br>ALA344,<br>LEU441 | PRO268,<br>VAL310,<br>HIS358,<br>ARG359,<br>ALA361 |
| Spike | ACE2 | TYR473,<br>ALA475,<br>GLN493,<br>GLY496,<br>GLN498,<br>THR500,<br>ASN501,<br>GLY502 | SER19,<br>GLU23,<br>ASP38,<br>ASN330,<br>LYS353,<br>GLY354,<br>ASP355 | GLU484 | LYS31 | TYR453,<br>LEU455,<br>PHE486,<br>TYR489,<br>TYR505 | LYS31,<br>HIS34,<br>MET82,<br>LYS353 |
| ACE2 | FP | GLU564 | HIS291 | TYR215 | SER255 | LEU91,<br>PRO565 | HIS291 |

**Table 3:** Comparative docking score and binding free energy of docked complexes of FP, Spike and ACE2.

| Complex | Zdock score | Binding free energy (kcal/mol) |
| --- | --- | --- |
| Spike & FP | 21.52 | -3867.90 |
| Spike & ACE2 | 16.40 | -1315.89 |
| Spike-FP & ACE2 | 18.58 | -1653.68 |

## Discussion

The COVID-19 pandemic has resulted in the loss of millions of lives and continues to pose significant financial burdens with rise of novel superspreading mutants having immune-evasive potential (26). Therefore, gaining a comprehensive understanding of the pathogenesis of SARS-CoV-2 remains crucial for a better managing the infection and preparing for future outbreaks. Extensive evidence emerged during the pandemic that highlighted the involvement of the complement system, particularly the dysregulation of the alternative pathway in COVID-19 pathogenesis and its immunopathological consequences (11, 27, 28). Additionally, it has been observed that the spike protein of SARS-CoV-2 can directly activate the alternative pathway (13). Out of the two key regulators of the alternative pathway, a reduction of FH levels and elevated levels of FP in severe SARS-CoV-2 infection and in those with fatal outcomes have been reported (29, 30). However, the role of FH and FP in initial immune response against SARS-CoV-2 infection independently of complement activation, remains unexplored. Thus, in this study, we investigated FH and FP immune functions in SARS-CoV-2 inflection. This study demonstrates that FH and FP, acting as soluble PRRs of the innate immunity, can directly interact with both SARS-CoV-2 spike and RBD proteins and modulate the SARS-CoV-2 infectivity. A549 cells expressing human ACE2 and TMPRSS2 (A549-hACE2+TMPRSS2 cells) were used as a relevant in vitro respiratory epithelium model (10). SARS-CoV-2 viral pseudoparticles were utilised as a safe model to study SARS-CoV-2 infection (10).

FH serves as a critical negative regulator in the complement alternative pathway (6). Notably, lung fibroblasts have been found to secrete FH locally, emphasising its role in lung homeostasis and immune regulation (31). On the other hand, FP acts as a positive regulator of the alternative pathway, with infiltrating neutrophils being a primary local source of FP secretion via secretory granules within the lung (7). Interestingly, both FH and FP have been shown to bind and inhibit influenza A virus (IAV) cell entry, their inhibitory effects on IAV infection are dependent on the specific IAV subtype (6, 7). Additionally, FH and FP have been demonstrated to attenuate inflammatory responses in A549 cells, further highlighting their immunomodulatory properties (6, 7).

Cell binding assay showed that FH significantly reduced lentiviral pseudoparticle binding to and infection of A549-hACE2+TMPRSS2 cells, whereas FP and TSR4+5 enhanced cell binding and infectivity. Thus, we conducted in-silico studies, which revealed that FP interacted with spike protein through TSR4+5 domains, forming a tripartite complex. This complex exhibited a high binding affinity to the host ACE2 receptor via TSR4, leading to an increased affinity between the virus and host ACE2 compared to the virus without FP. These observations suggest a possible mechanism by which FP enhances virus infection and replication, resulting in the poor prognosis observed in infected individuals. Luciferase reporter assay revealed that pre-treatment of SARS-CoV-2 lentiviral pseudoparticles with FH significantly reduced viral transduction within A549-hACE2+TMPRSS2 cells. Conversely, treatment with FP or TSR4+5 resulted in an increase in viral transduction compared to the control. Therefore, FH is a cell entry inhibitor for the SARS-CoV-2, whilst FP is a viral entry enhancer into lung epithelial-like cells. Anti-FP polyclonal antibodies seem to reverse FP-mediated enhancement of viral cell entry and binding.

Neutrophils serve as a reservoir for properdin and can release it rapidly via secretory granules upon activation (32). This local release of properdin from neutrophils is believed to be the primary factor influencing alternative pathway activity, as plasma properdin levels are typically low (32). Notably, neutrophils have been implicated in driving inflammatory responses during SARS-CoV-2 infection (33). The findings suggest that sequestering excessive FP may have the potential for an effective therapeutic strategy in limiting immunopathogenesis for severe SARS-CoV-2 infection.

Dysregulation of inflammatory response is one of the primary drivers in the transition of SARS-CoV-2 infection to moderate/severe COVID-19 (34, 35). Elevated serum levels of IL-1β, IL-6, IL-8, and TNF-α have been observed in SARS-CoV-2 infection, and are associated with the severity of the disease (34, 35). High levels of IL-6, TNF-α, MIP-2 and IL-8 gene expression have been found in alveolar type II cells challenged with SARS-CoV-2 (36). Furthermore, both mild and severe SARS-CoV-2 infections have been characterised by dysregulation of NF-κB activities (37–39). NF-κB signalling is vital for an effective immune response toward viral infection (40). Nevertheless, dysregulation of NF-κB functions is central to SARS-CoV-2 immunopathogenesis (37), because it has been associated with high levels of gene expression and proteins of pro-inflammatory mediators, including IL-1, IL-2, IL-6, IL-12, TNF-α, IL-8, MIP-1, MCP1 and RANTES in severe COVID-19 (37, 41–43), and hence, its therapeutic significance (44, 45). In this study, the gene expression levels of NF-κB were downregulated in A549-hACE2+TMPRSS2 cells challenged with FH-treated SARS-CoV-2 alphaviral pseudoparticles (FH-treated cells). In contrast, FP or TSR4+5 upregulated the NF-κB transcript expression. We also found a decrease in NF-κB activation in A549-hACE2+TMPRSS2 cells challenged with FH treated SARS-CoV-2 spike protein, whist there was an increase in NF-κB activation in FP-treated cells. Thus, FH may reduce hyperinflammation via the downregulation of NF-κB. On the contrary, FP can be exacerbating pro-inflammatory response in SARS-CoV-2 infection in the pulmonary tissues.

Another pro-inflammatory key player in the inflammatory response to viral infection is IL-1β (46). SARS-CoV-2 influences the activation of IL-1β, which can subsequently affect IL-6 and TNF-α (47, 48). SARS-CoV-2-induced cytokine storm is facilitated by IL-1β (49). Elevated levels of IL-1β in the peripheral blood and bronchoalveolar lavage fluid have been observed in patients with severe SARS-CoV-2 infection (50, 51). IL-1ß targeted therapy prevents SARS-CoV-2-induced cell death (52). Also, IL-1 receptor blocking in the early stage of the disease was found to be an effective treatment against respiratory failure, cytokine storm development, and hyperinflammation in COVID-19 patients (53). In this study, we observed that FH-treated cells demonstrated a reduction in the gene expression levels of IL-1β, while they were higher in FP and TSR4+5 treated cells. Elevated serum levels of TNF-α have been observed in severe SARS-CoV-2 infection (54, 55). TNF-α is a crucial immune player in limiting viral infections (56, 57). However, high levels of TNF-α contribute to lung damage and poor outcomes in severe COVID-19 patients (55). Combination of anti-TNF-α and anti-IFN-γ therapy in severe SARS-CoV-2 infection was able to reduce tissue damage and mortality (58). The mRNA levels of TNF-α in FH-treated cells were lower, whereas they were higher in FP and TSR4+5 treated cells. The findings imply FH could reduce TNF-α associated complications in SARS-CoV-2 infection, whilst FP may promote them.

IL-6 serum levels were higher in individuals with SARS-CoV-2-associated pneumonia, linked to the disease severity and mortality (59). Elevated levels of IL-6 are associated with a poor prognosis due to its contribution to inflammation and cytokine storm (59). Interestingly, COVID-19 patients, who are at risk of suffering cytokine storm, seem to respond well to tocilizumab, a monoclonal antibody that targets IL-6 receptors (59). Decreased mRNA levels of IL-6 by FH whilst its elevation in FP or TSR4+5 treated cells appear to suggest that FH might prevent the transition of SARS-CoV-2 infection to the severe form of the disease through limiting dysregulation of IL-6 levels, whereas FP may contribute to promoting IL-6 abnormality.

Type I IFN (IFN-α & IFN-β) is a major cytokine in localising and preventing viral infection via inducing the expression of interferon-stimulated genes (ISGs) (60). Elevated IFN-type 1 (IFN-1) levels could contribute to hyperinflammation in the progression to severe SARS-CoV-2 infection via various mechanisms (60). However, early studies have shown limited IFN-1 response in SARS-CoV-2 infection (60). Recent reports on the role of IFN-1 in the development of severe SARS-CoV-2 infection have surfaced (60). Also, a retrospective study revealed that the administration of IFN-α early reduced mortality, whereas its use in severe infection increased mortality and delayed recovery (61). Here, we showed that FH-treated cells exhibited a reduction in IFN-α gene expression levels, while in both FP and TSR4+5 treated cells, an elevation was observed with respect to the controls. Thus, FH could contribute to the immune regulatory mechanism in preventing unnecessary inflammation by limiting the action of IFN-α in immunopathogenesis in disease progression. At the same time, FP may be a co-factor in the disease pathology.

In individuals with severe SARS-CoV-2 infection, IL-8 is associated with significant neutrophil infiltration, respiratory failure, and acute kidney damage (62). IL-8 is a potent pro-inflammatory cytokine crucial for activating and recruiting neutrophil cells during inflammation (62). Severe SARS-CoV-2 patients are more likely to experience neutrophilia than mild disease patients (62). A prior anti-CXCL-8 therapy prevents the onset of severe lung injury (63). In this study, we show that FH treatment downregulated IL-8 mRNA level, whereas it was upregulated in FP and TSR4+5 treated cells. These findings suggest the possible function of FH in preventing lung injury by inhibiting IL-8 in SARS-CoV-2 infection, while FP may exacerbate it in a complement activation-independent manner. Interestingly, various studies have demonstrated that high levels of neutrophil infiltration in severe SARS-CoV-2 infection is correlated with poor clinical outcomes (64). Since it is known that neutrophils release properdin from specific granules (7), FP may promote a feedback loop in which more IL-8 is produced in SARS-CoV-2 infection, leading to more neutrophil infiltration. That can result in elevated levels of FP being secreted, worsening the infection.

Lastly, elevated serum levels of RANTES were observed in mild and severe SARS-CoV-2 patients with respect to healthy control (65). RANTES (CCL5) is a potent leucocyte chemoattractant that induces the migration of various immune cells, including T cells, natural killer cells, dendritic cells, monocytes, basophils, and eosinophils (66, 67). High level of RANTES is associated with acute renal failure and liver damage in individuals with severe COVID-19 (68). Targeting RANTES early on in viral infection may improve viral clearance and help localise the viral infection (30). In this study, we found that FH-treated cells exhibited reduced gene expression levels of RANTES, whereas mRNA levels of RANTES were elevated in FP and TSR4+5 treated cell compared to their respective controls. These results suggest an immune role for FH in limiting viral infection and enhancing viral clearance via the involvement via RANTES in SARS-CoV-2 infection, while FP may delay viral clearance.

A high level of FP in severe COVID-19 patients (29) may be explained by its role in promoting both infection and inflammation resulting in the disease severity. Besides, the insufficient levels of FH in severe SARS-CoV-2 may contribute to the disease progression. While this study provides valuable insights into the potential immune protective role of FH and the immunopathological role of FP in COVID-19, additional experiments using clinical isolates from different variants and lineages are essential to understand the infection dynamics comprehensively. Moreover, *in vivo*, studies are required to evaluate the impact of local FH and FP in the lung microenvironment and explore combination therapies to mitigate complications associated with SARS-CoV-2 infection.

In summary, this study has unveiled a novel finding that FH and FP can interact with the RBD of the SARS-CoV-2 spike protein in a complement-activation independent manner. This interaction of FH hinders the virus from binding to its cell surface receptors, resulting in a reduction in SARS-CoV-2 infection of A549 cells that express both human ACE2 and TMPRSS2, while FP enhances the viral infection. Furthermore, the presence of FH led to a decrease in the expression levels of proinflammatory cytokines and chemokines, including IL-1β, IL-8, IL-6, TNF-α, IFN-α, NF-κB, and RANTES whereas these pro-inflammatory mediators were increased in the case of FP. Thus, complement proteins can act as a direct protective mechanism against viral infections; however, they, as is the case with FP, may also contribute to the disease severity independent of their roles in complement activation. This study sheds light on a mechanism through which our innate immune system offers protection as well as plays a part in SARS-CoV-2 immunopathology.

---

**Figure S1:**
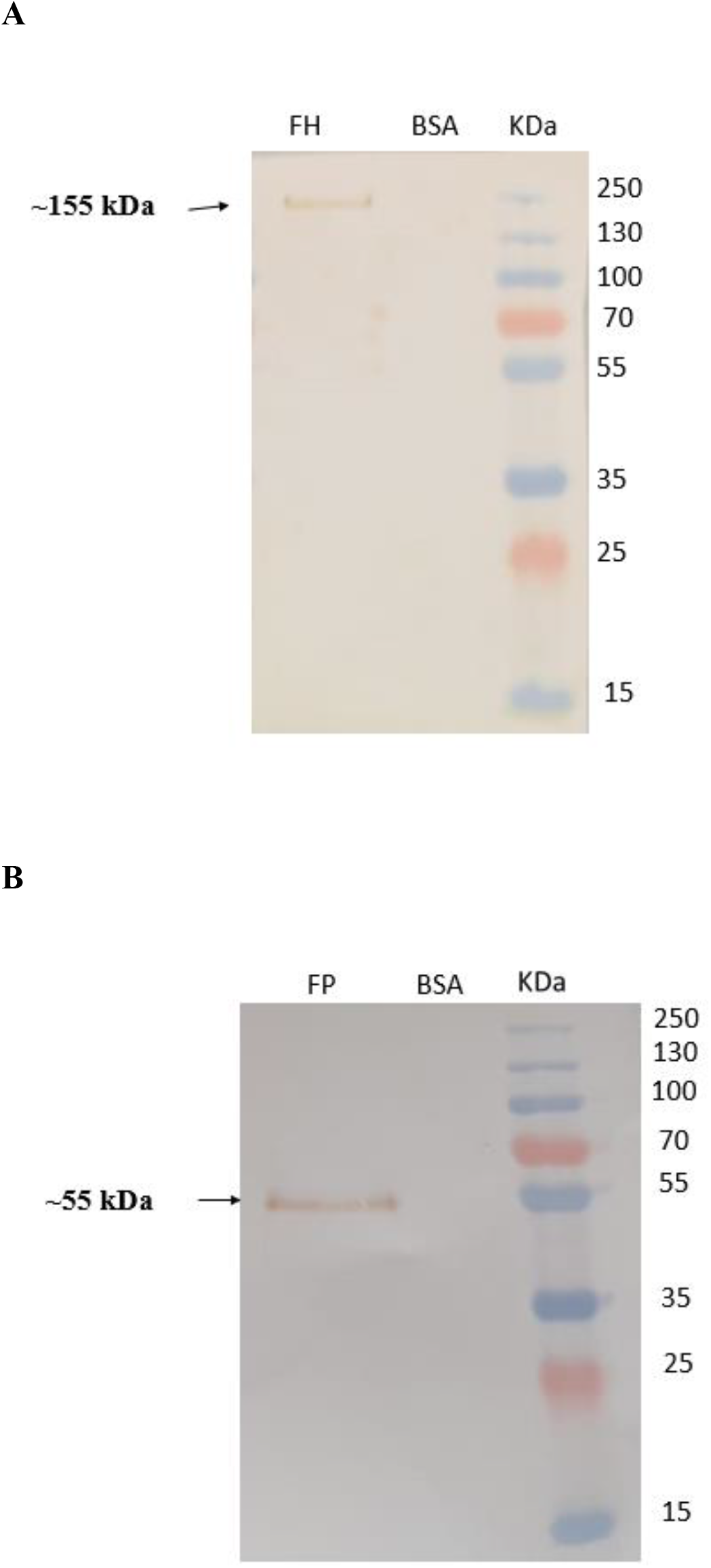
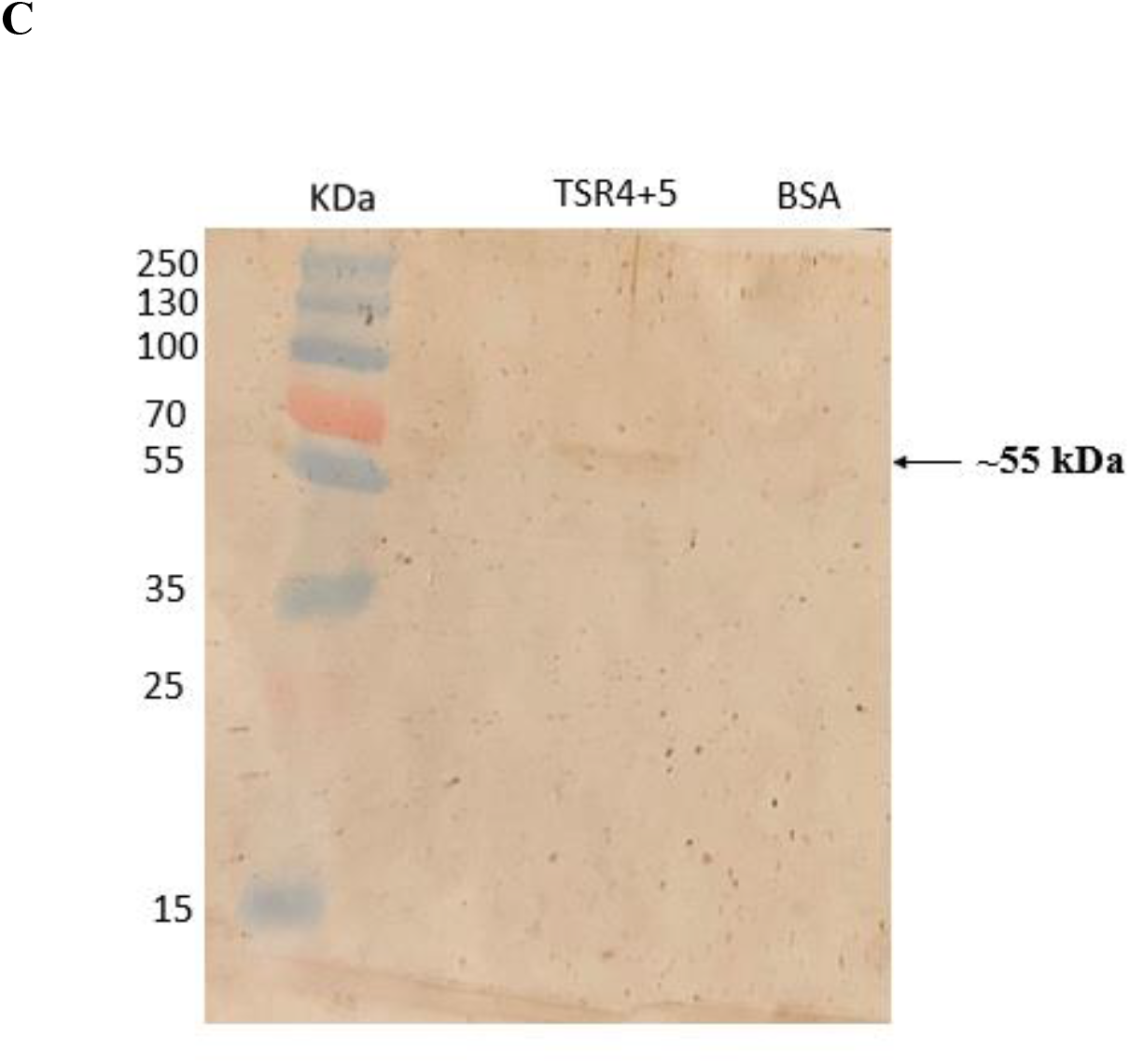
Characterization of purified complement proteins. The immunoreactivity of the purified complement proteins was assessed using western blotting. A PVDF membrane was employed and probed at room temperature for 1 h with specific antibodies (1:1,000 dilution). The antibodies used were rabbit-anti-human Properdin polyclonal antibodies, and mouse-anti-human FH monoclonal antibody (MRCOX23). Additionally, a protein ladder spanning a range of 250 to 10 kDa was included. Following the primary antibody incubation, the membrane was subsequently incubated with secondary antibodies, either goat anti-rabbit IgG HRP-conjugate or goat anti-mouse IgG HRP-conjugate (1:1,000 dilution), for 1 h at room temperature. The resulting bands corresponding to the respective proteins were observed. For Factor H was observed at ∼ 155 kDa (A), Properdin exhibited a band ∼ 55 kDa (B) and TSR4+5 was at ∼ 55 kDa (C), after developing the colour using 3,3′-diaminobenzidine (DAB) substrate.

**Figure S2:**
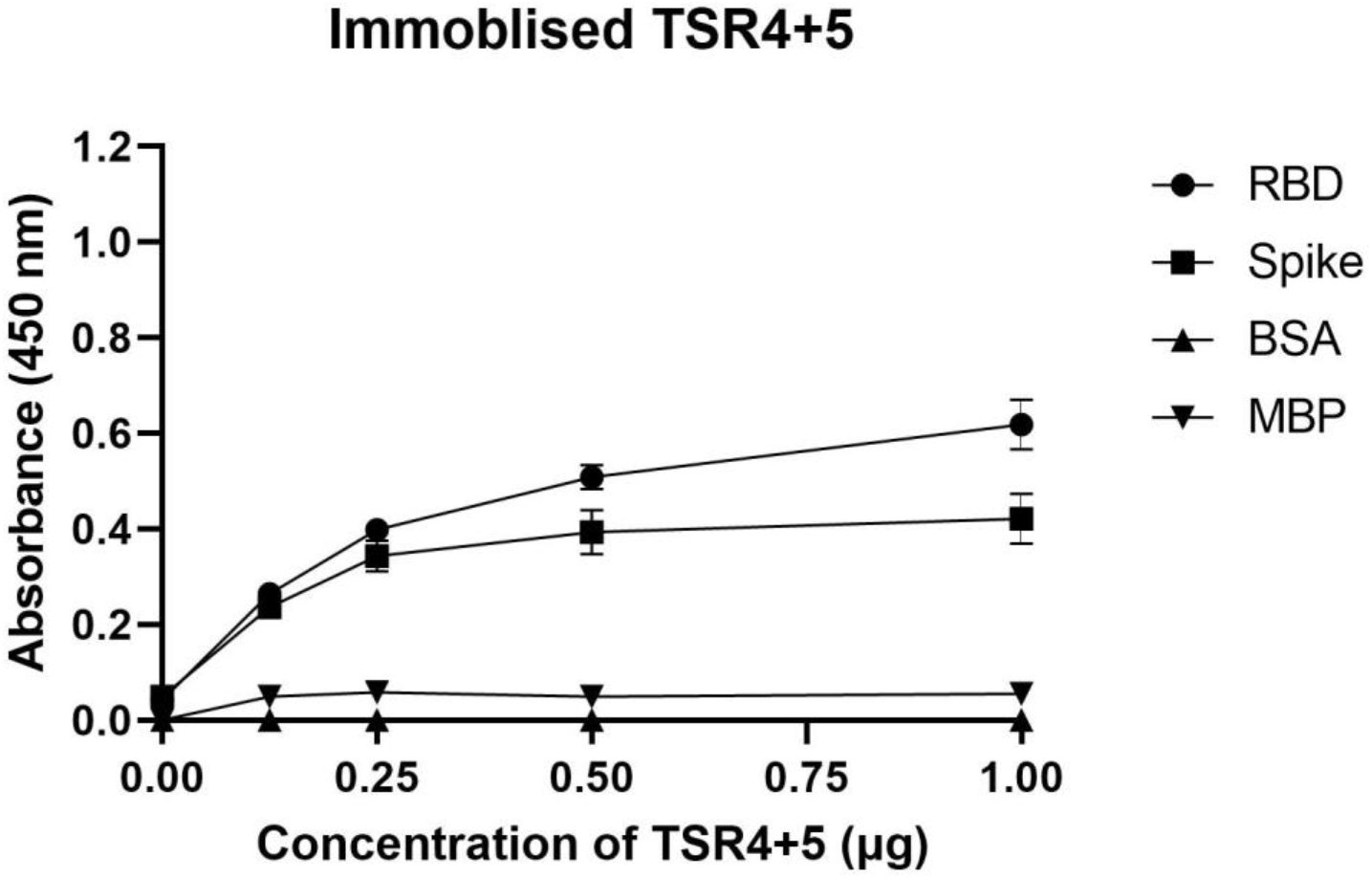
SARS-CoV-2 interacted with TSR4+5 via its S Protein RBD. TSR4+5 bound both SARS-CoV-2 Spike and RBD proteins in a dose-dependent manner. Decreasing concentration of immobilised TSR4+5 (1, 0.5, 1.25, and 0 μg/well) were coated in a 96-well plate using Carbonate-Bicarbonate (CBC) buffer, pH 9.6 at 4°C overnight. After washing out the excess CBC buffer with PBS, three times, a constant concentration of virus proteins (1 μg/well) was added to corresponding wells, followed by incubation at 37°C for 2h. After washing out the unbound proteins, the wells were probed with rabbit anti-SARS-CoV-2 Spike (1:5000; 100 μ/well). MBP and BSA were used as negative control.

**Figure S3:**
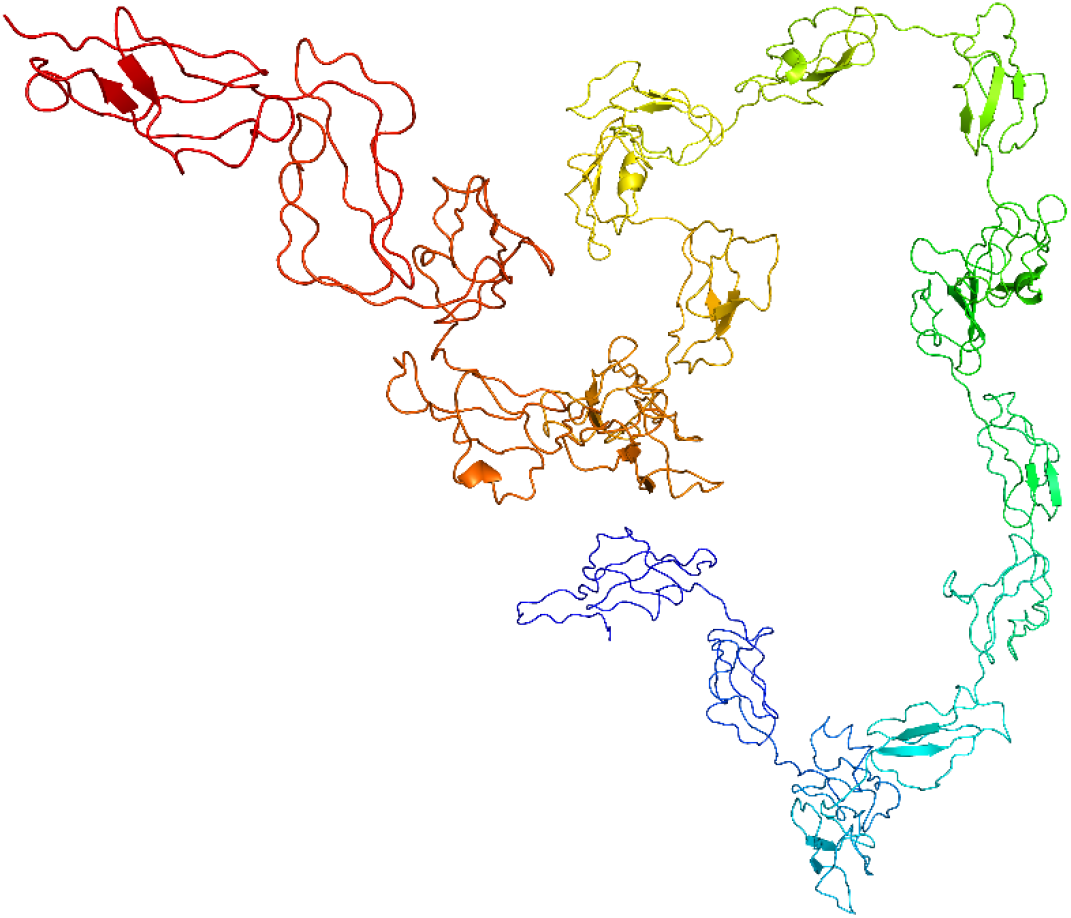
Cartoon representation of modelled FH structure. Full-length 3D structure of FH was modelled using Modeller10.1.

## Footnotes

1. Varghese, P. M.; Tsolaki, A. G.; Yasmin, H.; Shastri, A.; Ferluga, J.; Vatish, M.; Madan, T.; Kishore, U., Host-pathogen interaction in COVID-19: Pathogenesis, potential therapeutics and vaccination strategies. Immunobiology 2020, 225, (6), 152008.

